# The Bacterial Cytoskeleton Spatially Confines Functional Membrane Microdomains

**DOI:** 10.1101/2020.04.25.060970

**Authors:** Rabea Marie Wagner, Sagar U. Setru, Benjamin Machta, Ned S. Wingreen, Daniel Lopez

## Abstract

Cell membranes laterally segregate into microdomains enriched in certain lipids and scaffold proteins. Membrane microdomains modulate protein–protein interactions and are essential for cell polarity, signaling and membrane trafficking. How cells organize their membrane microdomains, and the physiological importance of these microdomains, is unknown. In eukaryotes, the cortical actin cytoskeleton is proposed to act like a fence, constraining the dynamics of membrane microdomains. Like their eukaryotic counterparts, bacterial cells have functional membrane microdomains (FMMs) that act as platforms for the efficient oligomerization of protein complexes. In this work, we used the model organism *Bacillus subtilis* to demonstrate that FMM organization and movement depend primarily on the interaction of FMM scaffold proteins with the domains’ protein cargo, rather than with domain lipids. Additionally, the MreB actin-like cytoskeletal network that underlies the bacterial membrane was found to frame areas of the membrane in which FMM mobility is concentrated. Variations in membrane fluidity did not affect FMM mobility whereas alterations in cell wall organization affected FMM mobility substantially. Interference with MreB organization alleviates FMM spatial confinement whereas, by contrast, inhibition of cell wall synthesis strengthens FMM confinement. The restriction of FMM lateral mobility by the submembranous actin-like cytoskeleton or the extracellular wall cytoskeleton appears to be a conserved mechanism in prokaryotic and eukaryotic cells for localizing functional protein complexes in specific membrane regions, thus contributing to the organization of cellular processes.

## INTRODUCTION

The organization of cell membranes influences all cellular processes and is essential for cell viability, yet our understanding of this organization remains poor. The original fluid mosaic model, which proposed that membrane proteins and lipids are distributed homogenously^1^, is only a rudimentary approximation of our current understanding of plasma membrane organization. Cell membranes are actually heterogeneous fluids, with different membrane lipid species laterally segregating into sub-micrometer domains, driven by their physico-chemical properties^2–4^. This heterogeneous organization of membrane lipids leads to a heterogeneous distribution of the embedded membrane proteins, which appears essential for their functionality^5^. Membrane microdomains are widely present, mobile structures that vary in size from transient protein clusters measured in nm^6^ to large micron-sized stable domains such as desmosomes^7^. These domains harbor proteins specialized in membrane trafficking, cell-cell adhesion, and signal transduction, and they provide targets for viral entry^8^.

It is now appreciated that membrane proteins can significantly impact membrane organization, in particular stabilizing membrane domains. For instance, the scaffold activity of the flotillin proteins FLO1 and FLO2 plays an important role in stabilizing some eukaryotic membrane rafts (“lipid rafts”)^9–11^; it would seem that this raft organization occurs in part through the capture and stabilization of specific lipids by flotillins^12,13^. In addition, the activity of the flotillin scaffold contributes to the recruitment of other raft-associated proteins, thus facilitating their interactions and correct functionality^14–16^. Other membrane-associated proteins, such as the actin-based membrane skeleton, are also thought to play an essential role in cell membrane organization, as detailed in the picket-fence model^17–19^. This model postulates that the membrane skeleton acts like a fence or corralling system, partitioning the cell membrane into small compartments. The direct anchoring of actin-associated membrane proteins (the pickets) to cytoskeletal filaments (the fences) beneath the cell membrane creates restrictive barriers that constrain the mobility of proteins and lipids to specific compartments or corrals. In plant cells, which are surrounded by a complex cell wall structure, membrane nanodomains are not only constrained by cortical cytoskeletal filaments but are also linked with the cell wall such that cell wall organization dramatically affects nanodomain mobility^20^. The interaction of plant flotillins (FLOTs) with the cell wall constituents is probably indirect, via interaction with cell-wall associated proteins that contain extracellular domains^20^. It is believed that this general mechanism of restricted diffusion of integral membrane proteins increases the probability of protein interaction within compartments. Events related to cell signaling or membrane trafficking may then be triggered upon protein interaction within compartments. In summary, the cellular cytoskeleton is now thought to play an important role in corralling biological processes within membrane microdomains^21^.

Cellular compartmentalization has traditionally been considered a fundamental step in the evolution of eukaryotic cell complexity. It was long thought that prokaryotes were too simple to require such sophisticated organization. Bacterial membranes are now known to laterally segregate certain lipid species into microdomains that accumulate at cell division sites or at cell poles^22–25^. This naturally leads to the heterogeneous distribution of many membrane proteins, especially those involved in cell division and cell-wall synthesis^26^. In addition, the lipids of prokaryotic membranes laterally segregate into functional membrane microdomains (FMMs) that compartmentalize proteins involved in different processes such as virulence, protein secretion, biofilm formation and antibiotic resistance^27–47^. The organization of FMMs in bacteria involves the biosynthesis and aggregation of isoprenoid-related membrane saccharolipids^47,48^ and their colocalization with flotillin-homolog proteins^47,49^. Bacterial flotillins, like their eukaryotic counterparts, are microdomain-associated scaffold proteins responsible for providing stability to FMMs and for recruiting protein cargo to the latter^27,32,50^. Thus, FMMs act as platforms for promoting the efficient oligomerization of protein complexes, and their organization influences cellular processes.

FMMs in the model bacterium *Bacillus subtilis* contain two flotillin-like scaffold proteins, FloA (331 aa) and FloT (509 aa), that homo- and hetero-oligomerize. FloA and FloT are distributed in discrete foci along the entire bacterial membrane, and display a highly mobile behaviour^49^. The correct spatial organization of FMMs likely prevents unwanted protein-protein interactions. How bacterial cells ensure that their FMMs are correctly distributed in the cellular membrane is poorly understood. Actin-like MreB and its homologs MreBH and Mbl are key components of the bacterial cytoskeleton and play an important role in bacterial membrane organization^51,52^. They form filaments underneath the membrane and interact with integral membrane proteins, such as the *mreB* operon partners MreCD and cell wall synthesis proteins such as RodZ, the penicillin-binding proteins (PBPs) PBPH, PBP2a and SEDS (shape, elongation, division and sporulation) proteins^53–57^. The MreB filaments and the associated cell wall synthesis machineries move circumferentially around the cell, perpendicular to the cell axis in a process driven by peptidoglycan synthesis^54–56,58^. The organization of MreB cortical filaments depends on the correct organization of the extracellular cell wall and *vice versa*. Thus, these are interconnected systems. In *B. subtilis*, the MreB filaments demarcate membrane regions with increased fluidity (RIFs)^51^, likely enriched in lipid II^59,60^. The bacterial actin-like cytoskeleton is essential for RIF formation as actin-homolog-deficient *B. subtilis* mutants are unable to generate RIF domains^51^. The bacterial actin-like cytoskeleton thus plays a crucial role in regulating membrane fluidity. It helps to maintain the correct organization of the membrane lipids and membrane proteins involved in cell wall synthesis and cell shape.

In the present work, we employ molecular, biochemical and fluorescence microscopy techniques to show that the *B. subtilis* cortical and cell-wall cytoskeleton provides a constraining barrier to FMM movement. This is akin to restrictions imposed on protein diffusion by the actin cytoskeleton and the cell wall in the eukaryotic membranes of plant cells^20^. Perturbations of the organization of the MreB filaments and of cell wall integrity interfere with the correct organization and mobility of FMMs. FMM mobility additionally depends on the size of the FMM and on its protein interaction partners, but not on treatments which alter membrane fluidity. The present results show that the bacterial cytoskeleton, including the cell wall and the MreB actin-like filaments, induce spatial confinement of the bacterial membrane – a mechanism that is conserved between eukaryotes and prokaryotes for the organization of membrane microdomains. The organization of bacterial cell membranes is thus more sophisticated than previously thought.

## RESULTS

### FloA and FloT assemblies show different lateral mobility patterns

The FMMs of *B. subtilis* contain two different flotillin-like scaffold proteins, FloA and FloT, that homo- and hetero-oligomerize. FloT is the larger protein (509 aa compared to 331 aa) and has an extended C-terminal region^31,32,61^ (Fig. 1a and Supp. Fig. S1). While *floA* is constitutively expressed, *floT* is upregulated by quorum sensing; its expression therefore increases during the stationary growth phase^32^. Fluorescence microscopy in projection mode showed strains labeled with FloA-GFP or FloT-GFP translational fusions to have approximately 12 visible FloA foci per cell, compared to approximately 6 larger FloT foci (Fig. 1b, c). These represent lower limits on the total number of foci, since domains at the top and bottom of cells would not be visible. Since the different subcellular organization of FloA and FloT might be indicative of different oligomerization patterns, the oligomerization of FloA and FloT was examined using Blue-Native PAGE (BN-PAGE). This technique allows the separation of membrane protein complexes in their natural oligomeric states^62^. BN-PAGE coupled to immunoblotting was used to identify membrane-associated protein complexes containing FloA or FloT. Distinct oligomerization profiles were detected for the FloA and FloT assemblies (Fig. 1d): the FloT assemblies were of higher molecular weight (MW), consistent with the larger FloT foci reported previously using super-resolution fluorescence microscopy^30,32^.

**Figure 1:**
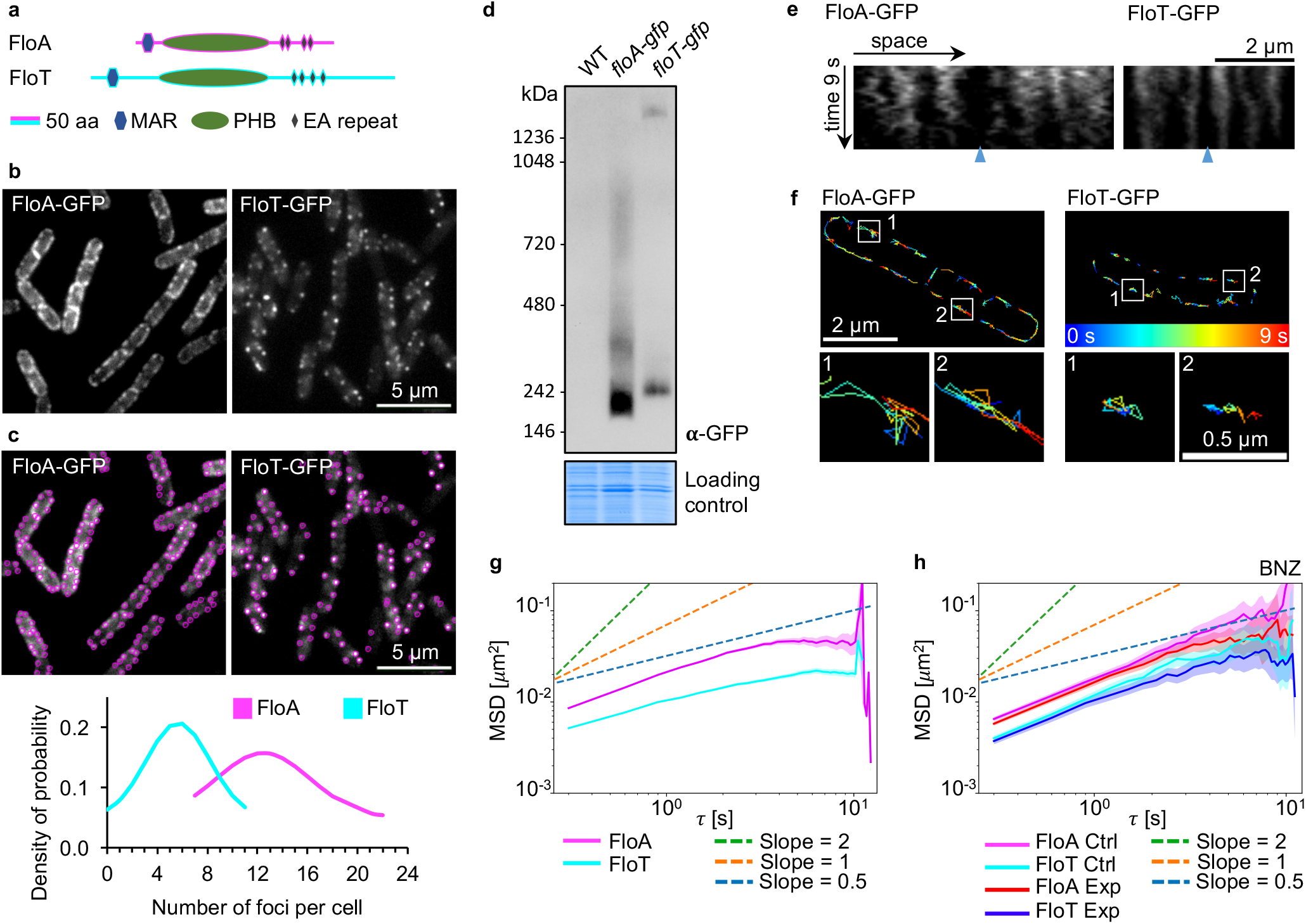
Flotillins diffuse with different mobilities. (Supplemental Figures S1-S4, Supplemental Table S1 and Supplemental Movies S1 and S2)**. a)** Models of FloA and FloT protein structure and domains. Flotillins are anchored to the membrane by the N-terminal membrane-anchoring region (MAR), followed by a prohibitin homologous domain (PHB), and a coiled-coil C-terminal region that contains glutamate-alanine repeats (EA). **b)** Fluorescence microscopy images showing the subcellular localization pattern of GFP-labeled FloA (left panel) and FloT (right panel) foci in *B. subtilis* cells. **c)** Spot detection exemplified with images of Fig 1b (top) and quantification of the relative abundance of visible fluorescence foci for FloA and FloT assemblies (bottom) in strains expressing FloA-GFP or FloT-GFP translational fusions (N=50 cells). **d)** Separation of membrane protein complexes in their natural oligomeric state by Blue-Native PAGE (BN-PAGE) coupled to immunoblotting, using GFP-labeled FloA- or FloT-expressing strains. A Coomassie-stained gel is shown as a protein loading control. **e)** Dynamic behavior of FloA-GFP (left) and FloT-GFP (right) represented in kymographs. The intensity of the signal in the membrane (*x*-axis) is represented over time (*y*-axis). Images were captured every 300 ms over 9 s. Cell poles of neighboring cells are marked with blue triangles. The membrane signal used for kymograph generation are specified by yellow arrows in Supp. Fig. S3a. **f)** Dynamic behavior of FloA-GFP (left) and FloT-GFP (right) represented by trajectories of flotillin foci followed over 9 s. Representative trajectories are shown in detail. Colors indicate elapsed time from blue=0 s to red=9 s. **g)** Plot of mean square displacement (MSD) against time of FloA-GFP (magenta) and FloT-GFP (cyan) (N≥13,988 trajectories from N=16 experiments). **h)** Plot showing the MSD analysis of FloA and FloT after increasing the membrane fluidity with benzyl alcohol. Ctrl=control condition, Exp=experimental condition (specified on the top), BNZ=benzyl alcohol (N≥399 trajectories). Plots show the averages with shaded area representing 95% bootstrap confidence intervals.

Flotillin assemblies are highly mobile. The movement of FloA and FloT assemblies was therefore monitored using time-lapse fluorescence microscopy; images were acquired every 300 ms over 9 s (Supp. Fig. S2a, b and Supp. Movie S1). The results were represented in kymographs, i.e., space-time plots of the membrane signal along the longitudinal axis of the cell over time (Fig. 1e and Supp. Figs. S2c, S3). In these kymographs, flotillin signals can be observed as continuous vertical tracks that move laterally within defined zones, pointing to a restriction to certain membrane areas. The FloA signal showed tracks with larger lateral displacements, while the FloT signal showed tracks with smaller lateral displacements, likely the result of more severe spatial confinement.

To gain more insight into the latter phenomenon, the trajectories of individual FloA and FloT membrane foci were visualized by analyzing the fluorescence signal using Fiji software equipped with the Trackmate plug-in (Supp. Fig. S2d). The FloA and FloT assemblies were associated with different types of trajectories. The FloA assemblies consistently showed both, movement over large membrane areas as well as spatially restricted movement. FloT assemblies largely showed spatially restricted movement with only rare long-distance migrations (Fig. 1f). Both FloA and FloT assemblies suffered obstruction to their trajectories, with lateral mobility often confined to a membrane region from which escape was apparently difficult. The most important difference between the trajectories of FloA and FloT assemblies lay in the frequency of escape from these transient obstructions. Fluorescence microscopy revealed the FloT assemblies to more often suffer temporary obstruction, whereas FloA rapidly moved from one obstacle to another. To further analyze these observations, we used the space and time information (*x*-, *y*- and *t*-positions) for the trajectories provided by Trackmate to plot mean square displacement (MSD) against time (Supp. Fig. 2e). In these plots, the *y*-intercept determines the level of movement and the slope characterizes the type of movement, with a slope of 1 corresponding to diffusion, a slope >1 to superdiffusion and a slope <1 corresponding to subdiffusion. MSD analysis of flotillin movement reveals a diffusive behavior for FloA showing a higher diffusion coefficient than FloT. FloT additionally shows subdiffusive behavior recognizable at large timescales (> 1 s) (Fig. 1g). Reduced mobility as visible for FloT is a characteristic of steric restriction. Overall, these results agree with the flotillin dynamics observed in the kymographs, confirming that FloA assemblies are more mobile than FloT assemblies, presumably due to increased restrictions imposed on FloT.

The influence of other factors related to the physico-chemical properties of the lipid bilayer, such as membrane fluidity, on the lateral mobility of FloA and FloT assemblies were also evaluated. Flotillins have been suggested to influence membrane fluidity^27,63^. It might therefore be that membrane fluidity regulates the lateral mobility of FloA and FloT assemblies. To test this, time-lapse fluorescence microscopy was used to analyze the mobility of the FloA and FloT assemblies when properties of the lipid bilayer including fluidity were altered. Cultures treated with benzyl alcohol (BNZ, 30 mM), which is known to increase membrane fluidity (Supp. Table S1)^51,64^, showed no significant differences in FloA or FloT diffusion coefficients (Fig. 1h; Supp. Fig. S4 and Supp. Movie S2). Thus, altering the lipid bilayer’s fluidity does not affect the mobility properties of the FloA and FloT assemblies and the hypothesis that flotillin membrane mobility is regulated by steric restrictions takes precedence.

### Flotillin mobility is determined by its cytoplasmic C-terminus

Both FloA and FloT have an N-terminal region that anchors them to the membrane adjacent to a prohibitin homologous domain (PHB, also known as SPFH [stomatin, prohibitin, flotillin and hflk/C]^65,66^) characteristic of the flotillin family (Fig. 1a). The results of experiments performed at our laboratory and others suggest that PHB domains associate with the cell membrane (via lipid binding) without traversing it^32,67^, as do other PHB-containing proteins such as podocin and stomatin^68,69^. The C-terminal region of flotillins, in contrast, is involved in protein-protein interactions and flotillin oligomerization^32,66,70–75^. We hypothesized that cellular components other than those involved in membrane fluidity affect the lateral mobility of FloA and FloT assemblies. Cellular components could exert a more pronounced effect on mobility either by membrane lipid interactions with the N-terminus of flotillin or by protein interactions with the C-terminus of flotillin. Distinguishing between these possibilities could reveal if either the N- or the C-terminus of flotillin controls its organization. To further explore this question, FloA and FloT variants with swapped N-terminal and C-terminal regions were genetically engineered. They were created to determine which protein region is associated with normal FloA or FloT subcellular distribution and movement. Since FloT is a larger protein with an extended C-terminus, it is this region that is the most obviously different between FloA and FloT. This C-terminus region is responsible for flotillin oligomerization^75^ and thus determines which interactions might occur among proteins. Accordingly, a chimeric version of FloA was generated that contained the C-terminal region of FloT (FloA_nt_T_ct_), as was a chimeric version of FloT that contained the C-terminal region of FloA (FloT_nt_A_ct_)^32^ (Fig. 2a). GFP-fused versions of these proteins were expressed under the control of the N-terminus flotillin promoter. The chimeric constructs behaved like wild type flotillins; they localized in the membrane, concentrating in a detergent-resistant membrane fraction (DRM - a subfraction of the membrane enriched in FMMs; in contrast, the detergent-sensitive membrane fraction, DSM, is enriched in membrane phospholipids)^9^ (Supp. Fig. S5a, b). The membrane was resolved by BN-PAGE to separate the oligomeric FloT_nt_A_ct_ and FloA_nt_T_ct_ species, and to compare them to the native flotillins by immunoblotting (Fig. 2b). The oligomerization profile of FloT_nt_A_ct_ was comparable to that of native FloA, showing assemblies of lower MW than FloT. In contrast, the FloA_nt_T_ct_ assemblies were comparable to native FloT assemblies, showing oligomeric species of higher MW (Fig. 2b). Thus, the C-terminal region is a key factor in this process and the N-terminal region would appear to play no vital role in maintaining the characteristic oligomerization profiles.

**Figure 2:**
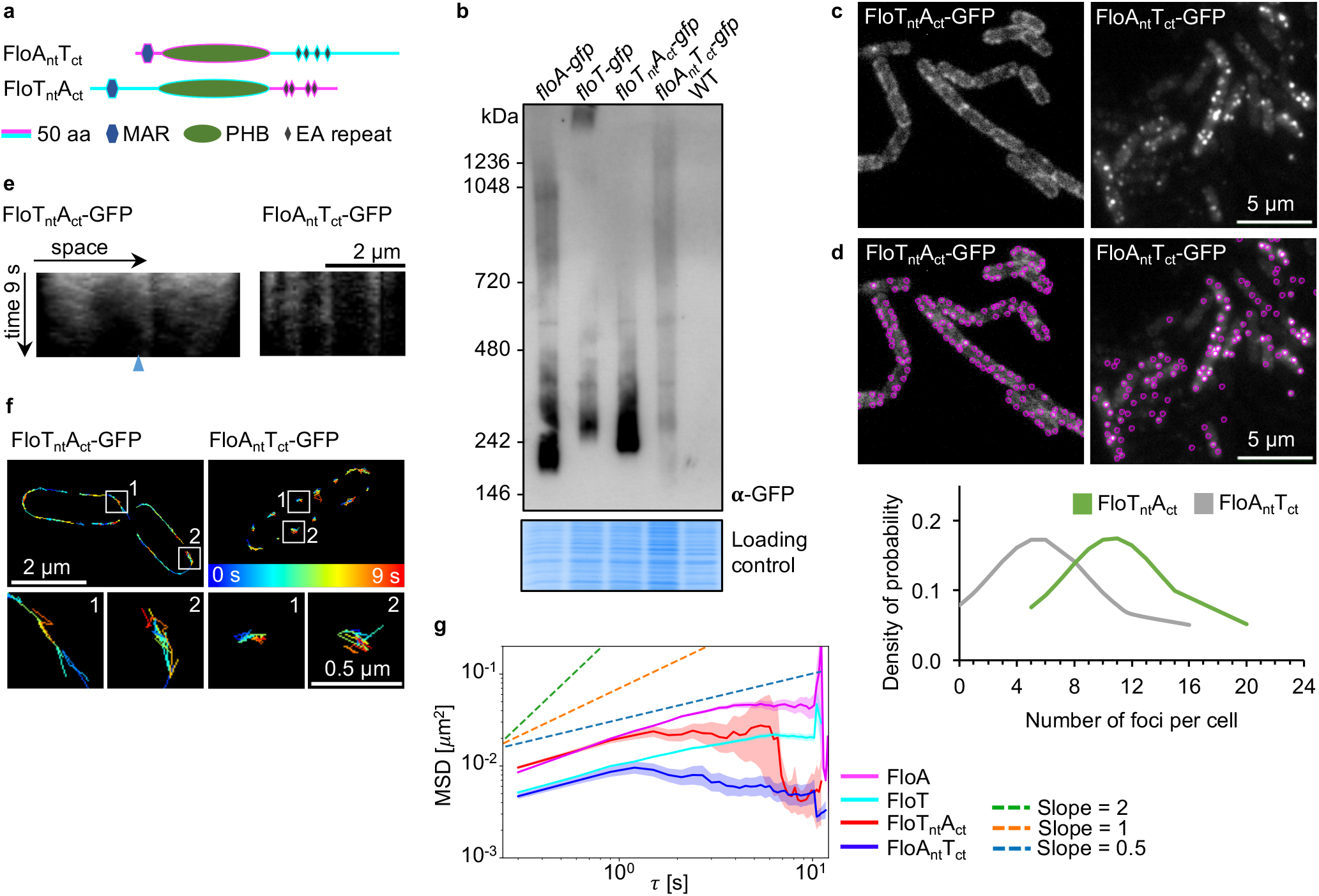
The C-terminus of flotillins determines their localization pattern and mobility. (Supplemental Figures S5 and Supplemental Movie S3)**. a)** Models showing the FloA_nt_T_ct_ and FloT_nt_A_ct_ chimeric flotillin protein structure and domains. **b)** Separation of membrane protein complexes in their natural oligomeric state using BN-PAGE coupled to immunoblotting, using strains expressing GFP-labeled FloA, FloT, FloT_nt_A_ct_ or FloA_nt_T_ct_. A Coomassie-stained gel is shown as a protein loading control. **c)** Fluorescence microscopy images showing the subcellular localization pattern of GFP-labeled FloT_nt_A_ct_ (left panel) and FloA_nt_T_ct_ (right panel) foci in *B. subtilis* cells. **d)** Spot detection exemplified with images of Fig 2c (top) and quantification of the relative abundance of fluorescence foci of GFP-labeled FloT_nt_A_ct_ and FloA_nt_T_ct_ assemblies (N=50 cells) (bottom). **e)** Dynamic behavior of FloT_nt_A_ct_-GFP (left) and FloA_nt_T_ct_-GFP (right) represented in kymographs; images were captured every 300 ms over 9 s. Cell poles of neighboring cells are marked with blue triangles. The membrane signal used for kymograph generation are specified by yellow arrows in Supp. Fig. S5c. **f)** Dynamic behavior of FloT_nt_A_ct_-GFP (left) and FloA_nt_T_ct_-GFP (right) represented in trajectories that flotillin foci followed over 9 s. Representative trajectories are shown in detail. Colors indicate elapsing time from blue=0 s to red=9 s. **g)** Plot showing the MSD analysis of FloA, FloT, FloT_nt_A_ct_ and FloA_nt_T_ct_. Plot shows the averages with shaded area representing 95% bootstrap confidence intervals (N≥946 trajectories).

The subcellular localization pattern of the chimeric flotillins was examined using fluorescence microscopy (Fig. 2c). The distribution pattern of FloT_nt_A_ct_ resembled that of FloA, showing an average of 11 small visible foci per cell, whereas the distribution pattern of FloA_nt_T_ct_ resembled that of FloT, organized into roughly 6 larger visible foci per cell (Fig. 2d). The typical subcellular localization pattern of each flotillin relied on its specific C-terminal region, in agreement with the above results.

The movement of the FloT_nt_A_ct_ and FloA_nt_T_ct_ assemblies was recorded by time-lapse fluorescence microscopy; images were acquired every 300 ms over 9 s (Supp. Movie S3) and the results represented in kymographs (Fig. 2e and Supp. Fig. S5c). The FloT_nt_A_ct_ signal showed continuous tracks with large lateral displacements, indicative of restricted movement hindered by recurrent obstructions. In contrast, the FloA_nt_T_ct_ kymographs showed tracks with smaller displacements, indicative of severe confinement of FloA_nt_T_ct_ movement (Fig. 2e). Compared to FloT_nt_A_ct_, the trajectories of the FloA_nt_T_ct_ chimeric variant suffered stronger temporary confinements, i.e., similar to the trajectories of native FloT. In contrast, FloT_nt_A_ct_ moved with more ease, escaping any obstructions more quickly - similar to the behavior seen for native FloA (Fig. 2f). MSD analysis revealed FloT_nt_A_ct_ to have a higher diffusion coefficient than FloA_nt_T_ct_ (Fig. 2g). Both corresponded to the diffusion coefficient of the respective C-terminal flotillin (Fig. 2g). Together, these results indicate that the C-terminal region of flotillins regulates flotillin movement in cell membranes. It is thus likely that different flotillin dynamics are the consequence of differences in their oligomeric states resulting from protein-protein interactions.

### Protein interaction partners influence flotillin dynamics

The FloA and FloT scaffold proteins form assemblies of different MW, in part because they tether distinct protein interaction partners. It was hypothesized that these partners might influence FloA and FloT dynamics. Flotillin-interacting proteins were therefore sought using pulldown assays in which a GFP-tagged version of FloA or FloT was expressed in a Δ*floA* Δ*floT* strain background. Flotillin-coeluted proteins were identified by mass spectrometry (Fig. 3a, Supp. Table S2). Enriched proteins were classified according to their functional groups (Fig. 3b). Thirty-four membrane proteins that interacted preferentially with FloA were identified, whereas 23 proteins were the preferential partners of FloT. Ten proteins showed non-preferential binding with FloA and FloT (Fig. 3c). A significant number of FloA-interacting protein partners that interacted with FloT_nt_A_ct_ was detected, whereas FloT-interacting protein partners interacted with FloA_nt_T_ct_ (Fig. 3d). All the proteins detected were represented in a heatmap and clustered according to their functional group^76^. This heatmap depicts the fold change protein abundance compared to the control strain (Δ*floA* Δ*floT* strain background expressing untagged GFP). The heatmap showed a similar behavior for all flotillin variants confirming that chimeric variants behave like the native flotillins. Mostly, only small differences in protein abundance can be detected between the samples. Nevertheless, close examination of the interacting protein’s abundance showed comparable binding levels for flotillin with its respective C-terminal chimeric variants (FloA + FloT_nt_A_ct_ and FloT + FloA_nt_T_ct_) (Fig. 3e and Supp. Table S2). Flotillin interaction partners were found in functional categories related to transport (FhuD, FeuA, RbsAB), signaling (RpoBC, SecY) and cell wall synthesis (PBP3, TagU, DltD, ErzA, MinD, FtsZ, MreC).

**Figure 3:**
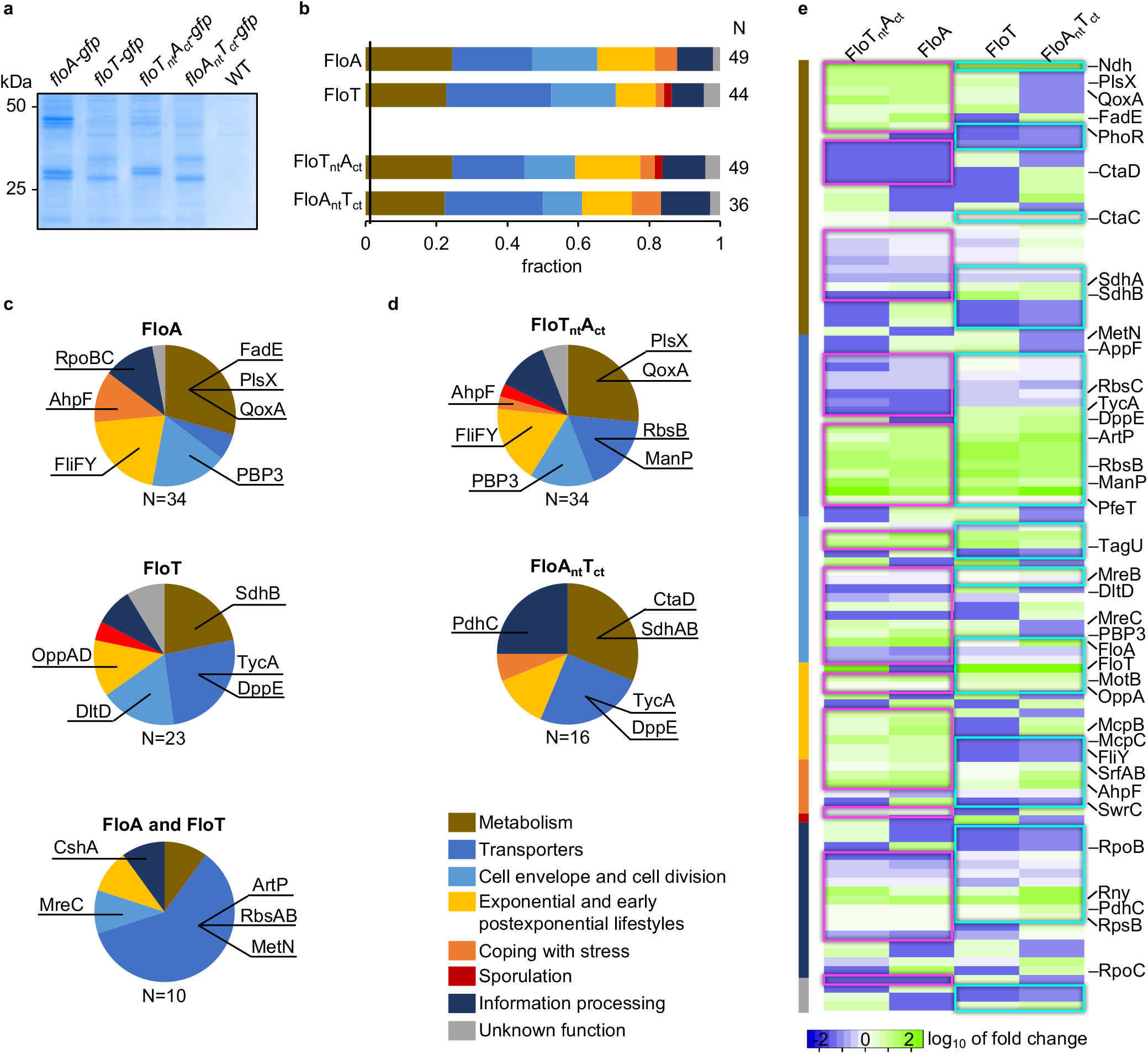
Flotillins interact with cell wall-related proteins. (Supplemental Table S2)**. a)** Coomassie staining of protein fractions coeluted with GFP-labeled flotillin variants. **b)** Column chart showing the fraction of proteins assigned to each functional category, using proteins that coeluted with the flotillin variants and that were enriched compared to the controls. **c)** Proteins that interacted with FloA (top) and FloT (middle) preferentially, or that showed no preferred interaction (bottom), were identified by pulldown assays followed by mass spectrometry analysis. Proteins are sorted according to their functional category and color-coded. **d)** Proteins that interact preferentially with FloT_nt_A_ct_ (top) and FloA_nt_T_ct_ (bottom), identified by pulldown assays followed by mass spectrometry analysis. Proteins are sorted according to their functional category and color-coded. **e)** Heatmap of coeluted proteins, categorized according to functional categories. Similarities between FloT_nt_A_ct_ + FloA and FloA_nt_T_ct_ + FloT are framed. Fold-change values are represented on the log_10_ scale. Some proteins are highlighted. Green = greater abundance than in control, blue = lesser abundance than in control. The heatmap was generated by Heatmapper software^102^.

Protein candidates that interacted with flotillin were selected to investigate whether the lateral mobility of flotillin assemblies is affected by their interacting partners. The penicillin-binding protein 3 (PBP3; encoded by *pbpC*) was detected in association with flotillins in pulldown assays (Fig. 3c and Supp. Table S2). The fluorescent probe Bocillin-FL, which binds specifically to PBPs^77^, was used to label PBPs prior to membrane fractionation to separate the DRM and DSM fractions. Using SDS-PAGE, PBP fluorescence signals were detected in association with the DRM and DSM fractions. Peptide mass fingerprinting was used for protein identification. PBP1 was identified in association with the DSM fraction, whereas PBP3 and PBP5 were mostly associated with the DRM fraction (Fig. 4a). PBP1 is part of the divisome and participates in lateral cell wall synthesis; PBP3 catalyzes transpeptidation and PBP5 is a carboxypeptidase^78,79^. GFP-PBP3 + FloA-mCherry and GFP-PBP3 + FloT-mCherry double-labeled strains were constructed and GFP-PBP3 used as bait to detect coelution with FloA or FloT in pulldown experiments. This revealed the preferential binding of PBP3 to FloA (Fig. 4b and Supp. Fig. S6). Total internal reflection fluorescence (TIRF)^80^ microscopy was used to detect PBP3 subcellular organization in membrane *puncta*, which showed a distribution pattern comparable to that of flotillins (Fig. 4c and Supp. Fig. S7a, b). In colocalization studies we detected higher levels of PBP3 signal associated with FloA signal, rather than FloT (Pearson correlation coefficient FloA-PBP3 r=0.657, FloT-PBP3 r=0.497) (Fig. 4d and Supp. Fig. S7d). In the absence of PBP3, using a Δ*pbpC* mutant (*pbpC* codes for PBP3), the dynamics of FloA and FloT assemblies only showed subtle changes (Fig. 4e; Supp. Fig. S8a, b and Supp. Movie S4). In counterpoint, the Δ*floA* mutants showed an increase in the PBP3 diffusion coefficient (Supp. Fig. S8 and Supp. Movie S5). Bocillin-FL-labelled DRM/DSM fractions of the Δ*floA* or Δ*floT* mutants were resolved in SDS-PAGE. Using this approach, an increase in PBP3 abundance was detected in the DRM fraction of the Δ*floA* mutant compared to WT whereas no differences were detected in the Δ*floT* mutant (Supp. Fig. S9). These results show that PBP3 mobility is reduced and its subcellular organization is affected by FloA.

**Figure 4:**
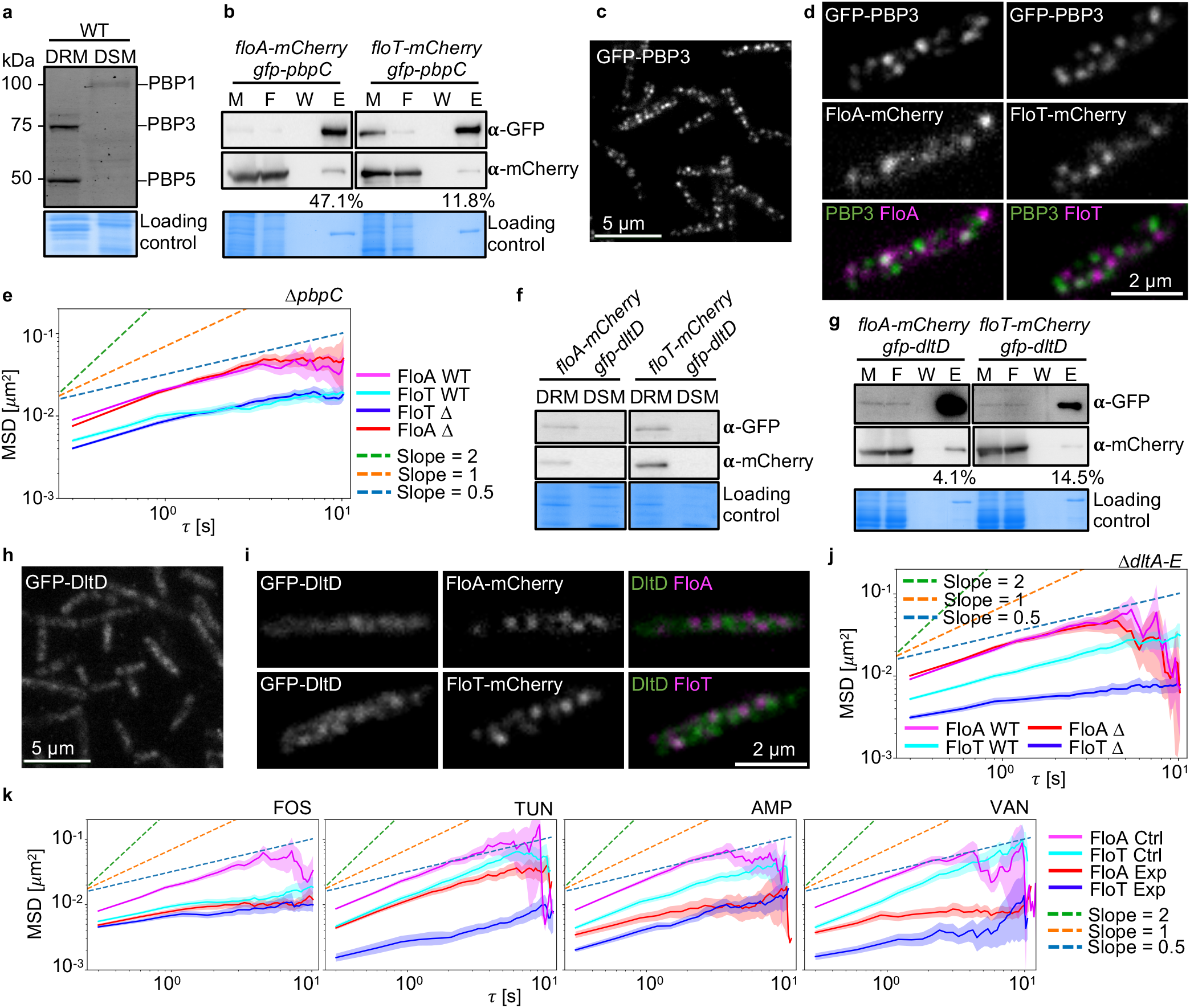
Interacting protein partners influence flotillin mobility. (Supplemental Figures S6-S14; Supplemental Tables S1 and Supplemental Movies S4-S6)**. a)** SDS-PAGE of DRM and DSM fractions of the WT. Bocillin-FL-labeled membranes were used to detect PBP membrane localization, and mass fingerprinting was used to identify PBPs. A Coomassie-stained gel is shown as a protein loading control. **b)** Immunodetection of FloA or FloT coeluted with PBP3 in GFP-PBP3 + FloA-mCherry or GFP-PBP3 + FloT-mCherry double-labeled strains. GFP-PBP3 was detected using α-GFP antibodies (left and right columns; upper lane). The mCherry signals for FloA (left column, lower lane) and FloT (right column, lower lane) were detected using α-mCherry antibodies. Numbers indicate relative mCherry elution band intensity compared to normalized GFP elution band intensity. M=whole membrane fraction, F=flowthrough, W=washed material, E=elution fraction. **c)** Fluorescence microscopy image of GFP-PBP3 subcellular localization pattern. **d)** Fluorescence microscopy images showing the colocalization signals of GFP-PBP3 + FloA-mCherry (left) and GFP-PBP3 *+* FloT-mCherry (right) double-labeled strains. **e)** Plot showing the MSD analysis of FloA and FloT in Δ*pbpC* (N≥1275 trajectories). **f)** Immunodetection of DltD, FloA and FloT signals in the DRM and DSM fractions of GFP-DltD + FloA-mCherry (left) or GFP-DltD + FloT-mCherry (right) double-labeled strains. **g)** Immunodetection of FloA or FloT coeluted with DltD in GFP-DltD + FloA-mCherry or GFP-DltD + FloT-mCherry double-labeled strains. **h)** Fluorescence microscopy image of GFP-DltD localization pattern. **i)** Fluorescence microscopy images showing the colocalization signals of GFP-DltD *+* FloA-mCherry (left) and GFP-DltD *+* FloT-mCherry (right) strains. **j)** Plot showing the MSD analysis of FloA and FloT in Δ*dltA-E* mutants (N≥1134 trajectories). **k)** Plots showing the MSD analysis of FloA and FloT after cell wall synthesis inhibition (N≥297 trajectories). Ctrl=control condition, Exp=experimental condition (specified on the top). FOS=fosfomycin, TUN=tunicamycin, AMP=ampicillin, VAN=vancomycin. For e), j) and k) plots show the averages with shaded area representing 95% bootstrap confidence intervals.

The membrane protein DltD was preferentially associated with FloT in the pulldown experiments (Fig. 3c and Supp. Table S2). DltD forms part of the DltABCDE membrane-associated protein machinery that adds D-alanyl residues to teichoic acids in the cell wall, thus attenuating negative-charge repulsion interactions^81–84^. The DltD signal mostly associated with the DRM fraction (Fig. 4f). Next, GFP-DltD + FloA- or FloT-mCherry double-labeled strains were generated in order to perform targeted pulldown analyses. GFP-DltD coeluted preferentially with FloT-mCherry, whereas a weaker interaction was detected with FloA-mCherry (Fig. 4g and Supp. Fig. S6). TIRF microscopy was then used to analyze flotillin and DltD spatial localization. GFP-DltD showed a diffuse, heterogeneous signal slightly more frequently associated with FloT than with FloA signal (Pearson correlation coefficient FloA-DltD r=0.62, FloT-DltD r=0.7) (Fig. 4h, i and Supp. Fig. S7c, e). Using fluorescence microscopy, the Δ*dltA-E* mutant showed no differences in FloA mobility, whereas a reduced diffusion coefficient of the FloT assemblies and a subdiffusive behavior at large timescales (> 1 s) were detected (Fig. 4j; Supp. Fig. S8a, b and Supp. Movie S4). Control experiments showed that changes in mobility are not caused by electrostatic differences in the cell wall, but by the absence of the Dlt proteins themselves (Supp. Fig. S10). The less diffusive behavior of FloT in the Δ*dltA-E* mutant, indicates that the DltA-E proteins promote FloT mobility. The DltD membrane diffusion coefficient was increased in the absence of FloT (Δ*floT* mutant), while the increase was less pronounced in the absence of FloA (Δ*floA* mutant) (Supp. Fig. S8e, g and Supp. Movie S5). These results show that DltD influences FloT movement and that DltD mobility is affected by FloT.

Having determined that different cell wall-related proteins interact differently with FloA and FloT, and influence their movement, experiments were performed to determine the effect of perturbing cell wall synthesis. The inhibition of cell wall synthesis restrains the motion of cell wall synthesis-related proteins^51,54–56,85^; thus, several antibiotics that inhibit different steps of cell wall synthesis were used to examine their effect on flotillin dynamics (Supp. Table S1). These antibiotics included fosfomycin (FOS; 2.5 mg/ml), which inhibits MurA, the catalyst in the first step of cell wall synthesis; tunicamycin (TUN; 2.5 μg/ml), which inhibits MraY and TagO, which are respectively involved in lipid II and wall teichoic acid (WTA) synthesis^86,87^; and ampicillin (AMP; 1 mg/ml) and vancomycin (VAN; 5 μg/ml), which inhibit the last incorporation step of cell wall precursor into the existing cell wall^88^. The lateral mobility of the FloA and FloT assemblies was monitored during the inhibition of cell wall synthesis using time-lapse fluorescence microscopy every 300 ms over 9 s. Treatment with any of these antibiotics at concentrations that inhibited cell wall synthesis reduced the diffusion coefficients of FloA and FloT (Fig. 4k, Supp. Fig. S11 and Supp. Movie S6) and led to subdiffusive behavior in some cases, especially after VAN treatments. Inhibition of cell wall synthesis using these antibiotics did not affect bacterial viability and no correlation of flotillin mobility with differences in membrane lipid composition was observed (Supp. Figs. S12-S14). Thus, flotillin dynamics is affected by cell wall synthesis in *B. subtilis*.

### The cortical and cell-wall cytoskeleton confine flotillin movement

We investigated how cell wall synthesis affects flotillin dynamics. Cell wall synthesis drives the motion of MreB cytoskeletal filaments as well as the MreB-dependent regions of increased fluidity (RIFs) – key factors in organizing the distribution pattern and movement of many membrane proteins^51,59^. MreB cytoskeletal filaments interact with the cell wall synthesis machinery, which leads MreB filaments to move along the cell membrane reflecting the motion of peptidoglycan synthesis^54–56^. MreB filaments organize circumferentially around the cell, perpendicular to the long axis of the cell and orient along the greatest curvature of the membrane. Thus, peptidoglycan is synthesized circumferentially^85,89^. The loss of any component of the cell wall synthesis machinery impacts the motion of the remaining components. Thus, the loss or depolymerization of MreB interferes with cell wall synthesis and causes the cell to grow as a sphere^90–92^. Conversely, the loss of the cell wall causes the disorganization of the MreB filaments (they adopt random orientations), resulting in a lack of oriented motion within the membrane of spherical cells^85,89^.

A connection between the bacterial actin-like cytoskeleton and FMM movement was inferred from the pulldown experiment, which points to an association of the cytoskeleton-related protein MreC (operon partner of MreB) with FloA and FloT (Fig. 3c and Supp. Table S2). However, colocalization experiments of FloA/FloT and MreB actin-like filaments, using FloA-mCherry + GFP-MreB or FloT-mCherry + GFP-MreB double-labeled strains, showed no colocalization (Fig. 5a and Supp. Fig. S15). Instead, TIRF microscopy indicated mutually exclusive localization. Accordingly, the MreB actin-like cytoskeleton and FMMs did not co-occur in membrane fractionation experiments. DRM and DSM fractions were collected and immunodetection studies performed. FloA and FloT signals localized preferentially in the DRM fraction, whereas the MreB signal was mostly associated with the DSM fraction (Fig. 5b). Taken together, the results of these experiments show that the MreB actin-like filaments are spatially organized in membrane areas that exclude FMMs.

**Figure 5:**
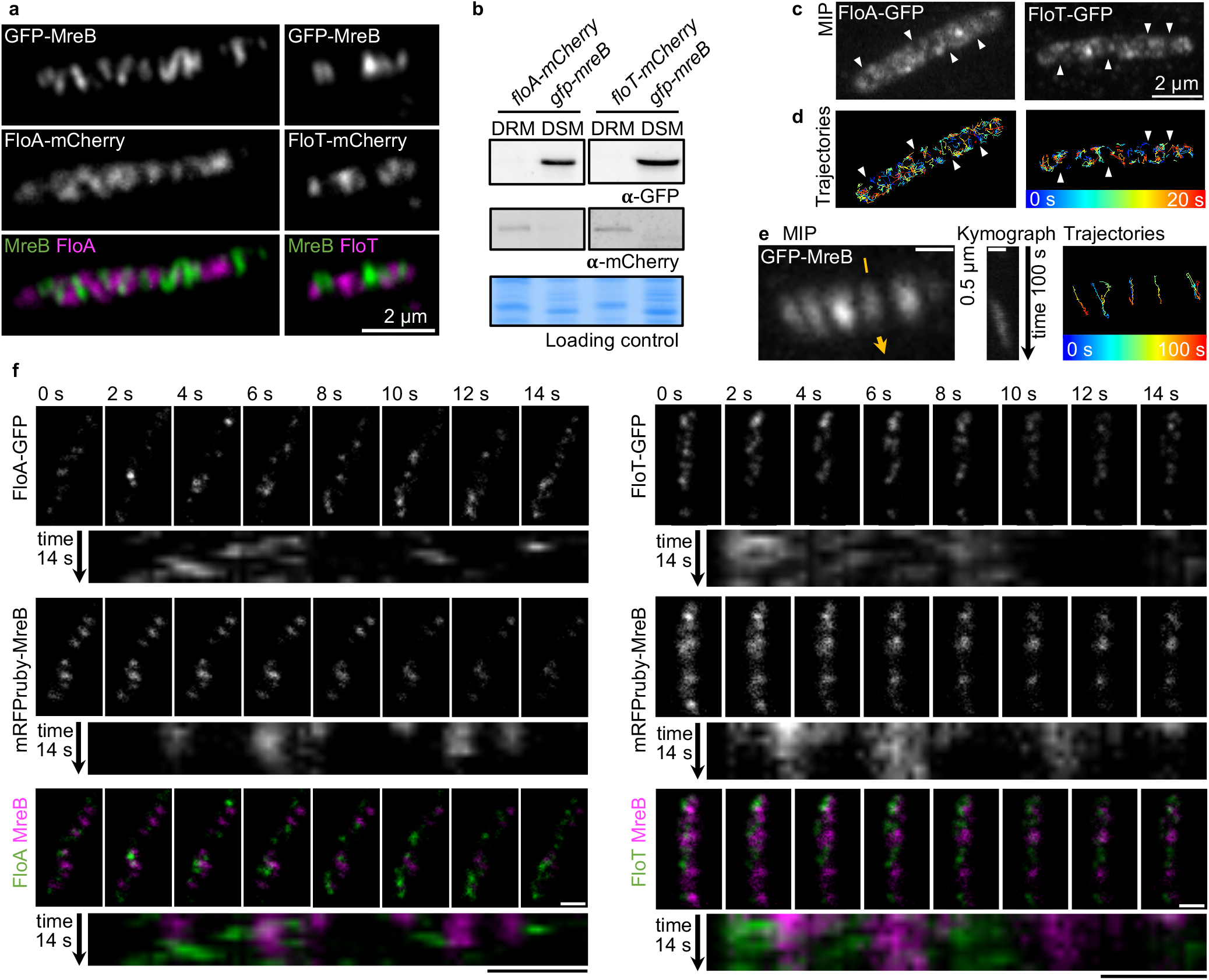
Flotillins and the actin-like cytoskeleton are spatially delimited. (Supplemental Figures S15-S17 and Supplemental Movies S7 and S8)**. a)** Colocalization of GFP-MreB with FloA-mCherry (left) and FloT-mCherry (right), as detected by TIRF microscopy. **b)** DRM and DSM fractions from GFP-MreB + FloA-mCherry (left) and GFP-MreB + FloT-mCherry (right) labeled strains. **c)** Maximum image projection (MIP) of FloA (top) and FloT (bottom) obtained from time-lapse TIRF microscopy sequences. Images were captured every 200 ms over 20 s. Arrowheads highlight membrane areas of no flotillin detection. **d)** Trajectories of FloA-GFP (top) and FloT-GFP (bottom) as created from time-lapse TIRF microscopy images. Arrowheads highlight the membrane areas without flotillin coverage. Colors indicate elapsed time from blue=0 s to red=20 s. **e)** GFP-MreB dynamics as revealed by TIRF microscopy in 1 s intervals over 100 s. MIP (left), kymograph (middle) of the signal indicated with the yellow arrow, and trajectories (right) of a representative cell are shown. Colors indicate elapsing time from blue=0 s to red=100 s. **f)** Time sequences of FloA-GFP (left) or FloT-GFP (right) and mRFPruby-MreB double-labeled strains, acquired with simultaneous TIRF microscopy (images captured every 2 s over 14 s) and kymographs corresponding to the signal of the longitudinal axis of the cell. Scale bars represent 1 μm.

It was shown above that the lateral mobility of FloA and FloT assemblies is constrained to membrane regions from which escape is difficult. The size as well as protein interaction partners that constitute these FloA or FloT assemblies influence the likelihood of escape. Larger FloT assemblies showed longer temporary confinement than smaller FloA assemblies. Based on these results, it was hypothesized that MreB cytoskeletal filaments (and MreB homologs) generate restrictive barriers in the bacterial membrane that confine the lateral mobility of FMMs. Since MreB organizes into filaments, it is likely that smaller FloA assemblies move more efficiently in the space between MreB filaments, slipping from one confinement to another more easily than larger FloT assemblies. To test this hypothesis, TIRF microscopy was used to monitor the dynamics of FMMs in relation to the localization of MreB filaments. To visualize the membrane area that flotillins covered during image acquisition every 200 ms over 20 s, maximal intensity projection (MIP) images were generated and the trajectories of both FloA and FloT examined. Membrane areas were detected in which FloA and FloT lateral mobility was concentrated; these alternated with membrane areas in which no flotillins were detected (Fig. 5c, d; Supp. Fig. S16 and Supp. Movie S7). The trajectories of the MreB filaments showed a movement chiefly in one direction perpendicular to the longitudinal axis of the cell^54,55,85,89^; their movement was much slower and more directed than that of the flotillin assemblies (Fig. 5e; Supp. Fig. S17a-c and Supp. Movie S8).

To gain a more detailed idea of the localization of FMMs in relation to MreB filaments, live-cell TIRF microscopy was performed with simultaneous image acquisition in the green and red channels. The localization of the GFP-labeled flotillins and mRFPruby-MreB in single cells was recorded every 2 s over a 14 s period. The difference in mobility between MreB and the flotillins precluded monitoring their movement simultaneously. The chosen time interval allowed the movement of MreB to be monitored whereas the FloA and FloT assemblies moved too rapidly to be properly followed (although their exact locations were mappable for each time point). The images show that FloA and FloT localized separately from MreB at all time points (Fig. 5f and Supp. Fig. S17d, e). In kymographs representing the signal along the longitudinal axis of the cell, the MreB filaments (magenta) showed steady lines consistent with their movement perpendicular to that axis. The FloA and FloT assemblies (green) appeared at different positions in the spaces between MreB filaments (Fig. 5f and Supp. Fig. S17d, e). Thus, the MreB filaments limited the motion of the flotillin assemblies.

FloA and FloT movement was monitored in cells that were unable to organize MreB filaments correctly. First, cells were limited for nutrients and oxygen, thereby abolishing formation of a sufficient membrane potential to maintain MreB functionality^93^. In these growing conditions, cells showed an increase in flotillin mobility (Supp. Fig. S18a-c). Second, to obtain more precise information on the influence of the MreB cytoskeleton on flotillin assemblies, a *B. subtilis* strain was examined that lacked the three actin-like protein homologs (MreB, MreBH and Mbl)^94^ and the differences in flotillin dynamics recorded. In this Δ*mreB* Δ*mreBH* Δ*mbl* strain, the loss of the actin-like cytoskeleton interfered with cell wall synthesis and caused cells to grow as spheres^90–92^ (Supp. Fig. S18d, e). It was hypothesized that flotillin lateral mobility should be increased in this strain if MreB-mediated spatial confinement really does influence flotillin dynamics. Further, the absence of any MreB confinement should equalize their diffusion coefficients (FloA assemblies move more efficiently between MreB filaments than larger FloT assemblies in standard growing conditions). To test this hypothesis, a GFP-FloA or GFP-FloT labeled Δ*mreB* Δ*mreBH* Δ*mbl* strain was designed and analyzed by fluorescence microscopy. The movement of FloA and FloT was recorded every 300 ms over 9 s. Kymographs revealed random movement and coverage over larger membrane areas with only occasional transient obstruction (Fig. 6a left panel; Supp. Fig. S18f-h and Supp. Movie S4). The trajectories of FloA and FloT showed spatially restricted movement coupled to periods of movement over large membrane areas (Fig. 6a right panel). MSD analysis revealed that the diffusion coefficient of FloT was increased, approaching that of FloA, whereas only a slight reduction in the diffusivity at early timescales (< 1 s) was visible for FloA (Fig. 6b). Overall, the loss of orientation or absence of actin-homolog filaments results in an increase in flotillin mobility and similar diffusion coefficients for FloA and FloT. Nonetheless, flotillin trajectories showed certain areas of confinement in cells lacking all actin-homolog filaments, which suggests that additional factors may influence flotillin movement.

**Figure 6:**
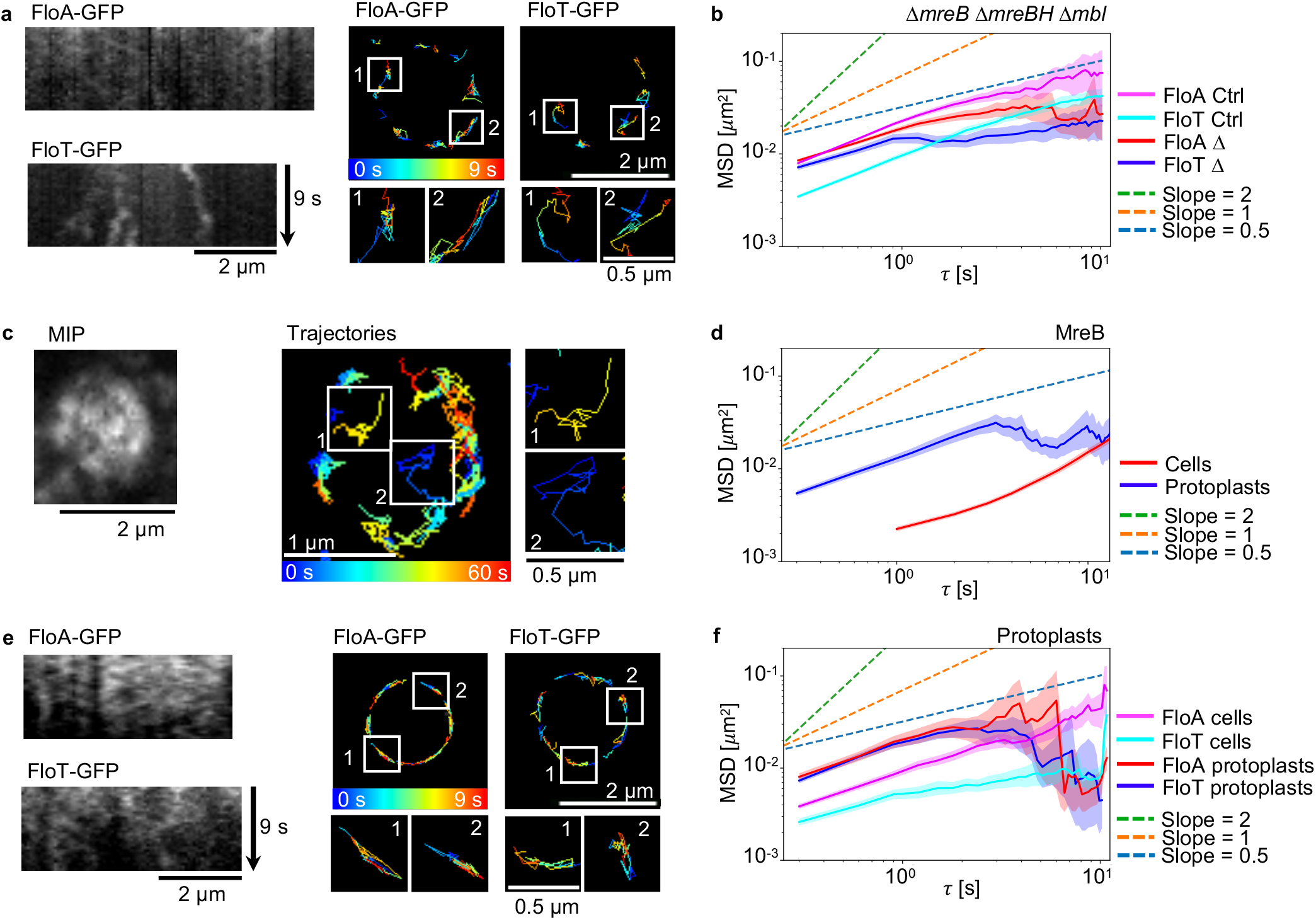
The actin-like cytoskeleton and the cell wall confine flotillin lateral movement. (Supplemental figure S18 and Supplemental Movies S4 and S8)**. a)** Dynamic behavior of FloA-GFP or FloT-GFP signal in Δ*mreB* Δ*mreBH* Δ*mbl* strain. Images were captured every 300 ms over 9 s. Kymographs (left) and trajectories (right) are shown. The membrane signal used for kymograph generation are specified by yellow arrows in Supp. Fig. S18f. Representative trajectories are shown in detail. Colors indicate elapsed time from blue=0 s to red=9 s. **b)** Plot showing the MSD analysis of FloA and FloT in the Δ*mreB* Δ*mreBH* Δ*mbl* mutant background strain (N≥666 trajectories). **c)** The cell wall of GFP-MreB labeled strains was digested in osmotically stabilizing conditions to generate protoplasts. The mobility of MreB was monitored with TIRF microscopy every 300 ms over 60 s. Results are shown in MIP (left) and trajectories (right). **d)** Plot showing the MSD analysis of GFP-labeled MreB in cells and protoplasts (N≥496 trajectories). **e)** Dynamic behavior of FloA-GFP or FloT-GFP in protoplasts. Images were captured every 300 ms over 9 s. Kymographs (left) and trajectories (right) are shown. The membrane signal used for kymograph generation are specified by yellow arrows in Supp. Fig. S18j. **f)** Plot showing the MSD analysis of FloA and FloT in protoplasts (N≥236 trajectories). For b), d) and f) plots show the averages with shaded area representing 95% bootstrap confidence intervals.

Our previous experiments demonstrated that cell wall organization also influences flotillin mobility, likely via interaction of flotillin with cell-wall associated protein partners. Thus, the bacterial cell wall could represent a supplemental cellular structure, in addition to the actin-homolog filaments, that influences flotillin mobility, similar to what occurs in eukaryotic plant cells^20^. To evaluate this hypothesis, we studied flotillin mobility in protoplasts. These are cells with the cell wall removed. Protoplasts have a spherical shape and still produce MreB filaments. However, the equal membrane curvature in all orientations of protoplasts causes MreB filaments to be disorganized and distributed randomly along the cellular membrane with a lack of oriented motion^85,89^. We monitored MreB in protoplasts and confirmed a random orientation of MreB filaments. Additionally, we detected a significant increase in MreB mobility in these conditions (Fig 6c, d; Supp. Fig. S18i and Supp. Movie S8). We hypothesized that if the cell wall restricts flotillin mobility, then protoplasts will show higher flotillin mobility than wild type cells. Hence, FloA-GFP- and FloT-GFP-labeled cells were treated with lysozyme and osmotically stabilized to render protoplasts^95^, and the movement of FloA and FloT assemblies monitored every 300 ms over a 9 s period by fluorescence microscopy^96^. The results showed a significant increase in the diffusion of FloA and FloT, with both reaching similar diffusion coefficients (Fig. 6e, f; Supp. Fig. S18j and Supp Movie S6). Their mobility patterns were comparable. Kymographs for the FloA signal showed random tracks with large lateral displacements and fewer membrane areas restricting FloA motion. The kymographs for the FloT signal were similar, revealing continuous, random movement with limited impediments to mobility, and covering large areas of the membrane. The FloA trajectories covered large areas of the membrane with apparently few transient obstructions. The trajectories of the FloT assemblies covered most of the membrane, quite similar to the FloA trajectories. These results suggest that the spatial confinement of FloA and FloT assemblies is much reduced in protoplasts; both flotillins moved faster and at comparable rates, irrespective of their oligomeric size or interaction partners. Altogether, our results demonstrate an interplay between the cortical and cell-wall cytoskeleton in restricting flotillin mobility, similar to what is observed in eukaryotic plant cells.

## DISCUSSION

In this work, we show that the organization and distribution of nanoscale functional membrane microdomains (FMMs) in bacterial cells is controlled by the cortical and cell-wall cytoskeleton, similar to what has been described for eukaryotic cells^20^. The movement of FMMs in the bacterium *B. subtilis* is determined by the interplay of several participating factors. The C-termini of FloA and FloT control protein-protein interactions and thus also dictate the sizes of these assemblies. FloA assemblies are smaller than FloT assemblies, and this size difference seems to play a role in the their subcellular organization and mobility. As flotillin mobility concentrates in membrane areas that are delimited by actin-like cytoskeletal filaments, which mainly affect larger flotillin assemblies, FloT mobility is more restricted. In addition, the bacterial cell wall plays a role in FMM mobility, likely via protein-protein interactions of the flotillin with cell-wall associated interaction partners. The interplay of these forces results in FloA assemblies moving faster than FloT assemblies.

The cell wall impacts flotillin mobility in two ways. First, upon antibiotic inhibition of cell wall synthesis, flotillin mobility is substantially decreased. The diffusion coefficient is reduced at short times and a subdiffusive behavior is visible on longer timescales. Possibly, diffusivity is reduced due to the general halt of cell wall synthesis machineries^54^ and the interaction partners such as PBP3 or DltD (Supp. Fig. S13a). This halt might also reinforce barriers that restrict movement due to an overall reduction of mobility in the membrane. In conclusion, the processes of cell wall synthesis stimulate flotillin mobility and might therefore actively drive flotillin motion. Second, in protoplasts, upon removal of the whole cell wall, the mobility of flotillins is increased. Furthermore, the mobility of FloA and FloT in protoplasts is the same, hinting towards an additional restricting mechanism that the cell wall applies to flotillins. As flotillins are located in the cytosolic side of the cell membrane, the connection to the extracellular cell wall is probably conferred by protein interaction partners. We saw that in protoplasts, MreB mobility is randomized and increased and the mobility of the flotillin interaction partners PBP3 and DltD is increased as well (Supp. Fig. S13b). Despite the evidence that the mobility of flotillins and of these cell wall interaction partners depend on each other, we observed low levels of colocalization and distinct mobility behaviors of all four proteins. Due to these differences, it is unlikely their interactions exist stably over long times. Rather, the interaction of flotillins with PBP3 and DltD is of transient nature, as has been reported for several other flotillin interaction partners^30^. Possibly, with the removal of the cell wall, the usual connections between the membrane and the cell wall are relieved from steric restraints. This general release from the friction of the cell wall might lead to increased mobility of flotillins and their equalization.

Together, the cell wall restricts flotillin mobility while at the same time its synthesis promotes motility, which might seem contradictory at first. On the one hand, flotillins are connected to the cell wall, likely by cell wall associated proteins, whose interaction with the cell wall restricts their mobility and thus also restricts flotillin mobility. Upon cell wall removal, this restriction is absent, and mobility is increased. On the other hand, while the cell wall is still present, inhibition of cell wall synthesis halts cell wall synthesis proteins, including those that connect flotillins to the cell wall, and thus flotillin mobility is reduced. Therefore, the cell wall exerts both positive and negative effects on flotillin mobility, likely mediated by transient interactions with cell wall associated proteins.

In addition to the cell wall, flotillin mobility – especially that of FloT – is restricted by the actin-like cytoskeleton. Removal of all actin-homologs releases FloT from spatial restrictions and increases its diffusion coefficient. Possibly, the differences between FloA and FloT in restriction by the actin-like cytoskeleton derive from the different sizes of FloA and FloT assemblies. Smaller FloA assemblies might circumvent MreB restrictions more easily, whereas FloT assemblies remain trapped inbetween MreB filaments for longer times. Our results are consistent with a model in which the lateral mobility of FloA and FloT assemblies occurs in membrane areas constrained both by MreB cytoskeletal filaments and by the cell wall, possibly via cell-wall associated interaction partners (Fig. 7). As a result, the diffusive behavior of FloA and FloT assemblies resembles the classical pattern observed in eukaryotic membranes: random diffusion of proteins over short timescales^97^ with reduced diffusivity or subdiffusion over longer timescales^52,98–100^. Notably, FloT diffusivity is lower than FloA on longer timescales. It remains challenging to discern the unique role of the cell wall in FMM mobility, as the organization and dynamics of the cortical cytoskeleton in bacteria, as well as of many membrane proteins, depend on the cell wall architecture (and *vice versa*). The loss of the cell wall causes the disorganization of MreB filaments^85,89^, and the loss of MreB filaments interferes with cell wall synthesis^90–92^. Thus, the MreB-mediated spatial confinement of cell membrane domains cannot be readily dissociated from the organization of the cell wall.

**Figure 7:**
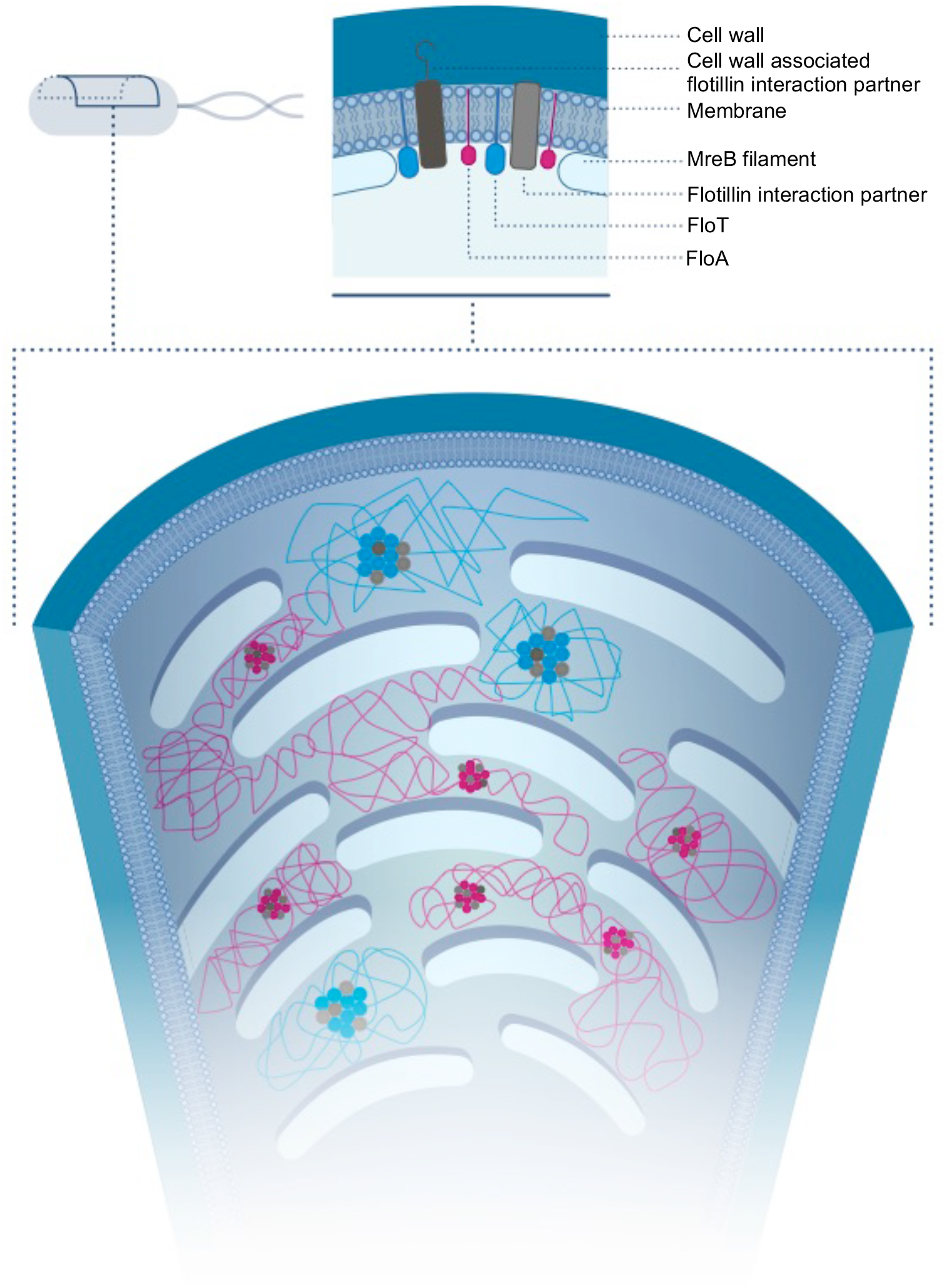
Model suggesting how MreB and its homologs and the cell wall sterically restrict the mobility of flotillin assemblies. The model depicts the membrane of *B. subtilis* seen from inside out. The MreB cytoskeleton filaments are localized on the inner leaflet of the membrane orientated perpendicular to the length axis of the cell. The cell wall is located on the other side of the membrane outside the cell. FloA (magenta) and FloT (cyan) interact with different protein partners some of which are linked to the cell wall. These assemblies move between the cytoskeletal filaments. The comparably slow movement of MreB does not impact flotillin dynamics and is not illustrated here. The bigger size of the assemblies of FloT as well as its protein-protein interaction partners prevent its long-distance diffusion restricted by MreB and its homologs. Furthermore, the diffusion of FloA and FloT is restricted by the cell wall possibly mediated by cell wall associated interaction partners.

The benefit of confined diffusion of FMMs in the bacterial membrane is unknown. This constraining mechanism may help distribute FMMs along the entire cellular membrane, by preventing FMMs from coalescing into one or a few large domains. Therefore, the cortical and cell-wall cytoskeleton may result in a more uniform dynamic coverage of the cells by FMMs, which could facilitate the efficient capture of FMM client proteins to promote their oligomerization. The efficient oligomerization of these complexes in general and their partitioning in particular might play an important role in bacterial processes like cell division, peptidoglycan synthesis, protein secretion, motility or signal transduction, which have been associated with the integrity of FMMs in previous studies^27,31,71,101^.

The FMM confinement strategy is comparable to that described for eukaryotic cells, thus implying that cellular mechanisms limiting the lateral movement of membrane nanodomains are conserved between prokaryotes and eukaryotes. The picket-fence model of the cell membrane of eukaryotic cells is currently the most widely accepted explanation for spatial confinement within biological membranes^21^. The model postulates that the cytoskeleton acts like a fence or corralling system, partitioning the cell membrane into small compartments, which constrain the diffusion of membrane proteins and lipids^18,19^. Thus, eukaryotic lipid rafts are confined in cytoskeleton-delimited membrane compartments, and the cytoskeleton provides an effective barrier against the large-scale coalescence of lipid rafts^21^. While bacterial cortical cytoskeletal restriction is broadly comparable to that described by the picket-fence model in eukaryotes, the bacterial cytoskeleton differs from the eukaryotic cytoskeleton. MreB forms independent filaments that are short compared to the actin filaments of eukaryotic cells, and as such do not form a well-defined meshwork underneath the plasma membrane. Despite these differences, the actin-like cytoskeleton still restricts flotillin mobility to some extent, but it cannot account for complete restriction of flotillin mobility.

In addition to the cortical cytoskeleton restrictive network, it has been recently shown that the movement of membrane nanodomains in plant cells is spatially restricted by the interactions of nanodomain-located proteins with the extracellular cell wall^20^. As a consequence, certain alterations of the cell wall increase the lateral mobility of plant flotillin proteins within the plasma membrane. The restriction of membrane nanodomain mobility by the cell wall is suggested to be indirect, likely via protein-protein interaction of plant flotillins with cell-wall associated proteins^20^. Indeed, flotillins in plants interact with diverse membrane proteins that contain extracellular domains, such as receptor-like kinases, transporters or AtTBL36, a protein that maintains the correct cellulose composition of the cell wall^20^. Similar to flotillin restriction in plant cells, we found the cell wall to account for additional limitations on flotillin mobility and especially for the differences in FloA and FloT mobility in our bacterial model *B. subtilis*. While the cytoskeletal organization differs, the cell wall of plants and bacteria is structurally and functionally comparable. Flotillins are located in the inner side of the membrane while connection with the extracellular cell wall is likely to be mediated by cell wall associated interaction partners, like PBP3 or DltD, akin to what has been proposed for plant cells^20^.

Our work provides evidence that prokaryotes and eukaryotes share fundamental mechanisms of membrane compartmentalization, presumably to maintain efficient protein-protein interactions, assist in the oligomerization of protein complexes for the optimal organization of cellular processes and metabolism, and to help avoid unfavorable protein stoichiometries and unwanted protein interactions. The cortical and cell-wall cytoskeletal structures act as effective barriers against the diffusion of membrane microdomains in both eukaryotes and prokaryotes^51,52^.

## ACKNOWLEDGMENTS

This work was funded by grant BFU2017-87873-P (MINECO, Spain) and ERC 335568 (European Researfch Council, EU), and in part by NIH Grants R01 GM082938 (N.S.W.) and NRSA 1F31CA236160 and training grant 5T32HG003284 (S.U.S.). We thank Prof. Jeff Errington (Newcastle University, UK) for providing the Δ*mreB* Δ*mreBH* Δ*mbl* Δ*rsgI* and GFP-PBP3-labeled strains, Prof. Peter Graumann (LOEWE-Zentrum für Synthetische Mikrobiologie, Germany) for sharing the GFP-MreB, -MreBH and -Mbl constructs, and Prof. Roland Wedlich-Söldner (University of Münster, Germany) for sharing the mRFPruby-MreB construct. The authors thank the fluorescence microscope facilities of the CNB-CSIC (Madrid, Spain) and the ZINF (University of Würzburg, Germany) for technical assistance and Scienseed (Madrid, Spain) for graphic design of Figure 7.

## MATERIALS AND METHODS

### Bacterial strains and culture conditions

The strains used in this study are derived from *B. subtilis* PY79 and recorded in Supplemental Table S4. *B. subtilis* cultures were grown in liquid LB (1% tryptone, 0.5% yeast extract, 0.5% NaCl) at 37°C or on minimal medium (MSgg: 5 mM potassium phosphate pH 7, 100 mM Mops pH 7, 2 mM MgCl_2_, 700 μM CaCl_2_, 50 μM MnCl_2,_ 50 μM FeCl_3_, 1 μM ZnCl_2_, 2 μM thiamine, 0.5% glycerol, 0.5% glutamate, 50 μg/ml tryptophan, 50 μg/ml phenylalanine)^103^ plates at 30°C. Media were supplemented with 0.5% xylose (exception MreB 0.01% xylose) and antibiotics if necessary (chloramphenicol 5 μg/ml, erythromycin 1 μg/ml, lincomycin 25 μg/ml, kanamycin 10 μg/ml, spectinomycin 100 μg/ml, tetracycline 5 μg/ml). *E. coli* cultures were grown in LB at 37°C supplemented with ampicillin 100 μg/ml if necessary.

### Construction of plasmids and strains

Mutants were constructed by PCR amplification of flanking regions (see primers in Supplemental Table S5), joining the individual fragments by overlap PCR using the Expand Long Template PCR System (Roche)^104^ and adding the PCR product to *B. subtilis* cells grown to late exponential phase in competence medium (100 mM potassium phosphate pH 7, 2% glucose, 0.1% casein hydrolysate, 0.2% potassium L-glutamate, 3 mM sodium citrate, 22 μg/ml ferric ammonium citrate, 3 mM MgSO_4_, 50 μg/ml of each L-tryptophan, L-phenylalanine and L-threonine)^105^. These conditions allow homologous recombination in cells and the selection of mutants by further plating the cultures with the appropriate antibiotics. The labeled actin-homolog mutation strain was constructed by successively deleting individual genes in flotillin-labeled strains by addition of genomic DNA of the Δ*mreB* Δ*mreBH* Δ*mbl* Δ*rsgI* donor strain (4277 of the Errington Lab^94^). The resulting strains were grown in LB supplemented with 20 mM MgSO_4_ and appropriate antibiotics. Plasmids containing labeled proteins were constructed by overlapping PCR and standard cloning techniques; final plasmids were transformed into competent *E. coli* and transformants selected in LB medium with ampicillin 100 μg/ml. Plasmids were then added to competent *B. subtilis* cells for transformation. Integrative plasmids were linearized before transformation. Transduction was performed with SPP1 phages in TY medium (LB supplemented with 10 mM MgSO_4_ and 100 μM MnSO_4_)^106^. To construct chimeric flotillins, flotillin domains were predicted using SMART^107,108^. The N-terminus was defined as the sequence reaching as far as the end of the predicted PHB domain. The rest of the protein was defined as the C-terminus. The N-terminus of one flotillin was joined with the C-terminus of the other by overlap PCR. Chimeric flotillins were expressed under the native promoter of the N-terminal flotillin at the *amyE* locus or from a replicative plasmid. To construct a useful replicative plasmid in *B. subtilis*, the backbone of the shuttle vector pJL-sar-gfp (which contains the ampicillin resistance cassette for selecting in *E. coli*, and the erythromycin resistance cassette for selecting in Gram-positive bacteria)^109^ was used, and the *sar-gfp* insert replaced with the multiple cloning site from pMAD^110^. Using this construct, different variants of the plasmid containing distinct antibiotic-resistance cassettes suitable for selection in Gram-positive bacteria were generated (Supp. Fig. S19). Plasmids and overlap PCRs of mutations were sequenced for confirmation. The strains and plasmids involved are listed in Supplemental Table S4.

### Fluorescence microscopy

Cells were grown to early (FloA) or late (FloT) exponential phase in liquid LB, washed with PBS and mounted on agarose-coated microscope slides for analysis at RT using a Leica DMI600B epifluorescence system (equipped with a Leica CRT6000 illumination system, a HCX PL APO oil immersion objective [100 × 1.47], and a Leica DFC630FX color camera) or an epifluorescence Leica DMi8 S System (equipped with a CoolLED pE4000 illumination system, a HCX PL APO oil immersion objective [100 × 1.47], and a Hamamatsu Orca-Flash 4.0 sCMOS camera) (for both microscopes: pixel size *x*=*y*=64.5 nm, image bit depth 16). Pictures were taken every 300 ms for at least 9 s with a 200 ms exposure time. If required, cells were previously treated with the following compounds for 90 min (if not stated differently): ampicillin 1 mg/ml, benzyl alcohol 30 mM (5 min), DMSO 2%, nisin 30 μM, tunicamycin 2.5 μg/ml or 0.025 μg/ml, valinomycin 60 μM (with Hepes 50 mM pH 7.5 and KCl 300 mM), or vancomycin 5 μg/ml. Protoplasts were generated by resuspending overnight cultures in SM buffer (0.5 M sucrose, 20 mM MgCl_2_, 10 mM potassium phosphate pH 6.8)^111^ with lysozyme (200 μg/ml) and incubating at 37°C for 30 min. Protoplasts were pelleted (4700 × g, 1 min, RT) and resuspended in SM buffer. An untreated control was included in every experiment and results of each experiment are presented with the respective control. After all treatments, flotillin mobility was analyzed and cell viability confirmed via CFU/ml counts. Additional control experiments were performed after specific treatments to check for possible changes in cell width, membrane permeability and potential, LTA abundance, fatty acid composition, and polar lipids. See below and supplemental material for details.

To study protein colocalization, cells were grown on MSgg plates for 24 h at 30°C. Biomass was fixed (2.5% paraformaldehyde, 0.03% glutaraldehyde, 10 mM sodium phosphate buffer pH 7.4; 10 min RT, 50 min on ice), washed three times with PBS, resuspended in GTE (50 mM glucose, 10 mM EDTA, 20 mM Tris-HCl pH 7.5), and stored at 4°C for maximum 1 week.

TIRF microscopy was performed using a Leica DMi8 S System (equipped with a TIRF Infinity HP module, a WSU unit with 488 nm and 561 nm solid state lasers [110 nm penetration depth], an HCX PL APO oil immersion objective [100 × 1.47], and a Hamamatsu Orca-Flash 4.0 sCMOS camera). Samples were prepared as previously described. To monitor the flotillin movement, images were acquired every 200 ms over a 20 s period. To monitor the MreB movement, images were acquired every 1 s over 100 s in cells and every 300 ms over 60 s in protoplasts. For simultaneous TIRF image acquisition, the microscope was additionally equipped with a W-VIEW GEMINI image splitter (Hamamatsu) with a dichroic mirror and bandpass filters specific for GFP (525/50) and mCherry (630/60). Samples were prepared as previously described and images acquired every 2 s for 14 s. The TIRF microscopy output consists of two-dimensional images of the curved surface of three-dimensional cells, and corrections to this spatial discrepancy have been used in several studies to quantify the diffusion coefficients of membrane proteins^52,112,113^. However, in the present work, the diffusion coefficients of the membrane proteins were compared under different experimental conditions without specifying absolute values, which obviated the need for any spatial correction of the TIRF images.

### Image processing

Microscopy images were processed using Fiji software^114^. For cell width measurements (N≥30 cells) the MicrobeJ plug-in was used^115^. Colocalization was analyzed using the JACoP plug-in^116^. Trajectories were determined using the Trackmate plug-in^117^, which detects spots and temporally (t-position) and spatially (x- and y-position) links them into individual trajectories according to certain input conditions (LOG detector 0.3 μm diameter spot detection [automatic spot quality filter, manually adjusted to include all visible foci, if necessary], LAP tracker [0.2 μm frame-to-frame linking, excluding gap closing, including splitting and merging], ≥ 4 foci per track; N=400 tracks) (Supp. Fig. S2). The output parameters of Trackmate depend on the input parameters (listed above). We therefore present results in comparison to their respective control and refrain from using absolute numbers. The trajectories r(t) = (x(t),y(t)) generated via Trackmate were used to calculate MSD as a function of time intervals *τ*, MSD(*τ*) = <[r(t+ *τ*)-r(t)]^2>, where the mean is over time t. MSD was calculated using code written in Python. MSD was calculated using trajectories (total N is specified for every experiment) collected from several independent experiments.

### Membrane harvesting

Cells were grown in liquid culture inoculated 1:100 from overnight cultures. Cell pellets were resuspended in PBS supplemented with 1 mM PMSF and lysed with lysozyme and sonication. After removal of any non-lysed cells, the whole cell extract was ultracentrifuged (200,000 × g, 1 h, 4°C) to harvest the membrane fraction. Membrane pellets were resuspended in PBS supplemented with 1 mM PMSF with or without DDM (n-dodecyl-B-D-maltoside (Glycon), the concentration for each experiment is specified) and homogenized in a sonication water bath.

### Western blot analysis and immunodetection

Protein samples were denatured by adding 1x SDS sample buffer and boiling for 5 min. Proteins were separated by 12% SDS-PAGE and stained with Coomassie or transferred to nitrocellulose membranes using the Trans-Blot Turbo Transfer System (Bio-Rad). Membranes were blocked in 10% skimmed milk in TBS-Tween 0.05% (TBS-T), incubated with the primary antibody for 2 h at RT and then with the secondary antibody for 1 h at RT. Antibodies were used at the following dilutions: rabbit-anti-GFP (monoclonal, Invitrogen) 1:1000 for native blots; rabbit-anti-GFP (polyclonal, Takara) 1:5000 for denatured blots; rabbit-anti-mCherry (polyclonal, BioVision) 1:5000; goat-anti-rabbit (BioRad) 1:20.000.

### Blue-native PAGE

The NativePAGE™ Novex system (Invitrogen) was used for native PAGE^62,118^. Cells were grown on MSgg plates prior to membrane harvest. Membranes were incubated with 0.1% DDM in 1x NativePAGE™ Sample Buffer with shaking at 4°C overnight. Unsolubilized membrane was pelleted, and solubilized membrane samples stained with 0.5% G-250 Sample Additive. Samples were run in 3-12% Bis-Tris gradient gels and transferred to PVDF membranes in a wet blot system (BioRad). Membranes were dried, the Coomassie removed with methanol to visualize the marker, and then further processed for regular immunoblotting.

### Pulldown assays and sample preparation for protein identification

A Δ*floA* Δ*floT* double knock-out strain background with GFP-labeled flotillin was used in our global pulldown analysis. Double labeled strains were used for targeted pulldown analysis. 500 μl of membranes from stationary cells were solubilized overnight with 1% DDM. Unsolubilized membranes were pelleted (17,000 × g, 30 min, 4°C) and the supernatant incubated with 25 μl of equilibrated GFP-Trap resin (ChromoTek) for 2 h at 4°C. The flowthrough was collected and the resin washed three times before elution with 50 μl of 2x SDS-loading buffer. In our global pulldown analysis (Fig. 3), the protein content of each sample was analyzed by mass spectrometry. For this, elution fractions were subjected to SDS-PAGE and stained with Coomassie. Each lane was cut into several pieces, fixed with methanol:acetic acid:H_2_O (4:1:5) and the proteins identified. The results for each lane is the assembly of the results of the individual pieces examined. For targeted pulldown analysis (Fig. 4b and h) the elution fractions were subjected to SDS-PAGE and western blot with immunodetection. Quantification was performed by normalizing the GFP signal intensity of the elution fraction and determining each mCherry signal intensity in relation to the GFP signal intensity.

### Protein identification

Excised protein bands were processed automatically using a Proteineer DP device (Bruker Daltonics). The bands were subjected to in-gel digestion and the peptides analyzed using a 1D-nano liquid chromatography apparatus coupled to a high-speed time-of-flight mass spectrometer with a nanospray III ionization source (Eksigent Technologies NanoLC Ultra 1D plus and TripleTOF 5600, SCIEX). Data were acquired with Analyst TF v.1.7 software and processed with PeakView v.2.2 software (both SCIEX). The detected peptides were compared against the genome of *B. subtilis* PY79 using the Mascot Server v.2.6.1 (Matrix Science).

For data analysis, only proteins with peptide-to-spectrum matches (PSM) of ≥2 were considered. These were normalized to the control (normalized spectral abundance factor, NSAF) and their fold change in abundance determined against the control. Membrane proteins were identified in the UniProt and NCBI databases via the provided accession numbers, and their functional categories assigned according to the *Subti*wiki^76^ classification. To further specify protein function, some functional categories were subdivided into subcategories (the subcategories ‘coping with stress’, ‘exponential and early post-exponential lifestyles’ and ‘sporulation’ belong to the main category “lifestyles”, and the subcategories ‘transporters’ and ‘cell envelope and cell division’ belong to the main category “cellular processes”). The binding of membrane proteins to FloA or FloT was deemed preferential when a threshold 1:1.25 fold enrichment difference was surpassed. Fold changes for membrane proteins were transformed to log_10_ values and visualized using Heatmapper software^102^ (input parameters: expression, no scaling, no clustering). Proteins not identified in a sample were manually assigned to the lowest fold-change value (shown in dark blue).

### Separation of detergent resistant DRM and sensitive DSM membrane fractions

The DRM and DSM fractions^119^ were separated using the CelLytic MEM Protein Extraction Kit (Sigma). 600 μl of lysis and separation buffer were added to 50 μl of resuspended membrane and incubated at 4°C overnight. Insolubilized membranes were pelleted (17,000 × g, 30 min, 4°C) and the supernatant incubated at 37°C for 20 min followed by centrifugation (3000 × g, 3 min, RT) for phase separation. The upper DSM fraction was removed, and the DRM fraction then washed three times with 400 μl of PBS (20 min on ice, 20 min at 37°C, centrifugation for phase separation). DRM and DSM were then precipitated with TCA, resuspended in 0.25 V of 1 x SDS sample buffer, and subjected to SDS-PAGE.

### Identification of penicillin binding proteins labeled with Bocillin-FL

Membranes were incubated with 100 μg/ml Bocillin-FL (Invitrogen) to label penicillin binding proteins (PBPs) for 30 min at 37°C prior to the addition of the lysis and separation buffer and subsequent DRM/DSM separation. Bocillin-FL was visualized in-gel with the ChemiDoc Flamingo Application (Bio-Rad). The Bocillin-FL signal does not persist after in-gel fixation or Coomassie staining. The PBPs of the Bocillin-FL bands were cut out under UV-light, fixed, and identified by peptide mass fingerprinting. Samples were in-gel digested and analyzed using MALDI-TOF/TOF (Abi 4800 MALDI TOF/TOF mass spectrometer, SCIEX). Data were acquired using ABi 4000 Series Explorer Spot Set Manager software and processed using ABi 4000 Series Explorer Software v.3.6 (both SCIEX). The detected peptides were compared against the genome of *B. subtilis* PY79 using the Mascot Server v.2.6.1 (Matrix Science). Only peptides with an individual score above the identity threshold were considered correctly identified, and attention paid only to identified PBPs.

#### Statistics

Sample size and error bars are specified for each experiment individually. Differences between samples and controls were examined using the unpaired two-sample Student’s t-test with Welch’s correction. Significance was set at p<0.05. Shaded regions for mean squared displacement curves represent 95% bootstrap confidence intervals.

#### Data availability

The microscopy data that support the findings of this study are available from the corresponding author upon request.

#### Code availability

Codes for MSD analysis are available from the corresponding author upon request.

