## Supplemental information for "The Bacterial Cytoskeleton Spatially Confines Functional Membrane Microdomains"

**This file contains:**

Supplemental Materials and Methods
Supplemental Figures S1 to S19
Supplemental Tables S1, S3 and S4
Supplemental Movie Captions S1 to S8
Supplemental References

**Additional Material in separate files:**

Supplemental Table S3
Supplemental Movies S1 to S8

### Supplemental Methods

#### Determination of antibiotic minimum inhibitory concentration

Minimum inhibitory concentrations (MICs) were determined in 96-well plates with serial dilutions of antibiotics in LB medium. Wells were inoculated with *B. subtilis* WT strains from overnight cultures to a final $OD_{600}$ of 0.05. Plates were incubated at 37°C for 20 h without agitation. MIC was determined in triplicate as the lowest antibiotic concentration that showed no bacterial growth.

#### Analyzing membrane properties with fluorescent dyes

Several dyes were used to visualize the activity of membrane-active compounds. The dyes were added for the last 15 min of the treatments. For the analysis of the membrane potential, 3,3'-diethyloxacarboxyanine iodide ($DiOC_2(3)$ (Sigma) was added to a final concentration of 30 nM. The cytoplasmic dye $DiOC_2(3)$ shifts from green to red fluorescence upon cell accumulation indicating unaltered membrane potential[1]. The LIVE/DEAD BacLight Bacterial Viability Kit (Invitrogen)[2], which contains the dyes SYTO-9 and propidium iodide, was used to determine differences in membrane permeabilization (dilution 1:1000 for both dyes). Propidium iodine (red fluorescence signal) only accumulates in the cytoplasm of cells with compromised membrane permeability.

#### Membrane lipid analysis

For cellular fatty acid composition and polar lipid analyses, cells were grown until exponential phase and treated with the compounds for 90 min. 200 mg of lyophilized cell pellet was used in analyses. To determine the fatty acid composition, the samples were subjected to gas chromatography coupled to mass spectrometry. Polar lipids were analyzed by 2D-TLC. Cellular fatty acid and polar lipid analyses were performed by the identification service of the DSMZ in Braunschweig, Germany.

#### Lipoteichoic acid (LTA) isolation and ELISA

We have used a variation of a previously reported protocol[3]. Cultures were harvested after antibiotic treatment, washed with 0.5 and 0.25 V of 100 mM sodium citrate pH 4.7, and resuspended in the same buffer (1 ml for $OD_{600}$=20). 700 µl sample was lysed in a GenoGrinder with 250 µl glass beads, the glass beads sedimented (200 x g, 5 min, RT), and the supernatant incubated with the same volume of n-butanol with agitation at RT for 30 min. The phases were separated (17,000 x g, 20 min, RT) and the lower aqueous phase containing LTA collected. A 100 µl sample was added to each well of a highbinding 96-well plate (Sarstedt) and incubated overnight at RT. The bound samples were blocked with 3% skimmed milk in TBS-T for 2 h and incubated with 50 µl 1:50 lipoteichoic acid monoclonal antibody (clone 55, HycultBiotech) for 2 h. Wells were washed 5x 3 min with TBS-T and incubated with 50 µl 1:500 HRP conjugated goat-anti-mouse antibody (Thermo Fisher) for 2 h followed by 5x 3 min washes with TBS-T. 100 µl of developing solution (50 mM sodium dihydrogen phosphate, 25 mM citric acid, 0.4 mg/ml O-phenylenediamine dihydrochloride [Sigma], 0.4 µg/ml hydrogen peroxide, pH 5) was added, incubated in the dark, and the reaction stopped after 20 min with the addition of 50 µl $H_2SO_4$ 1 M. The absorbance at 490 nm was then determined.

**Detergent resistance**

Cultures were harvested after antibiotic treatment, washed with PBS, and resuspended in 1 ml PBS. Dilution series were performed in 96-well plates and 100 µl culture added to PBS or PBS with SDS at a final concentration of 0.05%. After 15 min incubation time at RT, 10 µl samples were spotted on LB plates and incubated. Difference in survival between treatments with or without SDS were determined and compared to controls.

### Supplemental Figures

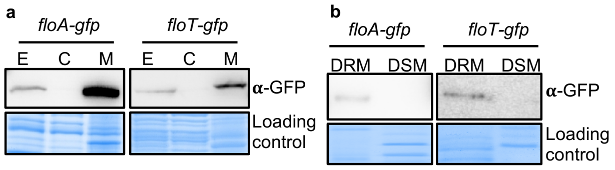


**Supplemental Figure S1: Flotillins localize in the detergent-resistant membrane (DRM) fraction of the membrane** (Related to main Figure 1)**. a)** Cell fractionation and immunodetection of GFP-labeled FloA (left) and FloT (right) membrane proteins. A Coomassie-stained gel is shown as a protein loading control. E=whole cell extract, C=cytosol, M=membrane. **b)** Fractionation of the membrane shows that FloA (left) and FloT (right) preferably localize in the DRM fraction.

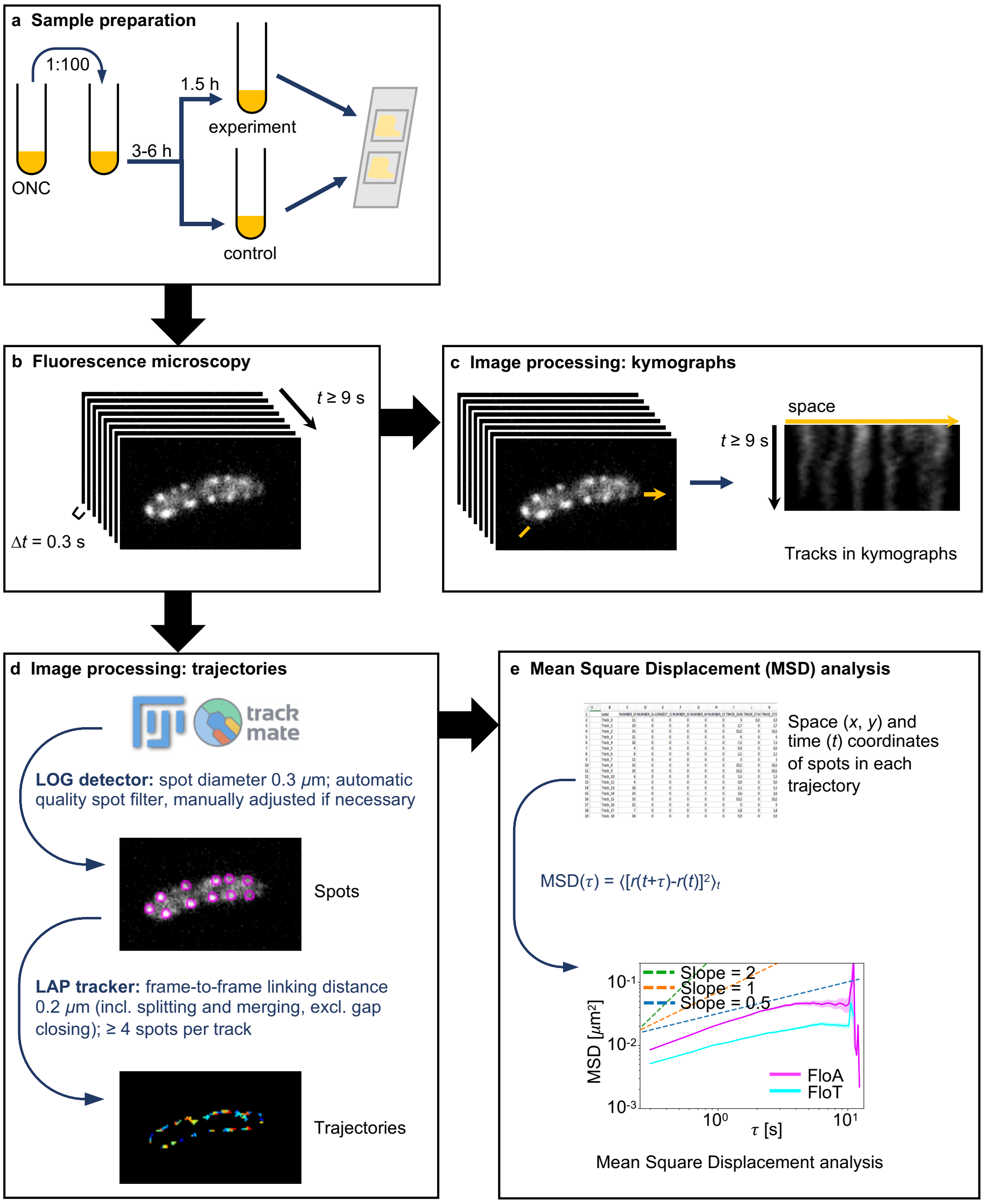


**Supplemental Figure S2: Experimental protocol for determining membrane foci mobility** (Related to main Figure 1)**. a)** Cultures were grown from overnight pre-cultures and allowed to grow until early (FloA) or late (FloT) exponential phase and subjected to experimental conditions. **b)** Experimental condition and control samples were subjected to fluorescence microscopy image acquisition (images were captured every 300 ms over 9 s). **c)** Images were processed to create kymographs by using the signal along the membrane indicated with a yellow arrow. **d)** Microscope images were analyzed using the Trackmate[4] plug-in for FIJI[5]. It detects individual foci and links them into trajectories according to the specified input parameters. The *x*-, *y*- and *t*-coordinates of each trajectory were recorded in each experiment. **e)** Mean square displacement analysis were conducted using *x*-, *y*- and *t*-coordinates of the spot positions along each trajectory.

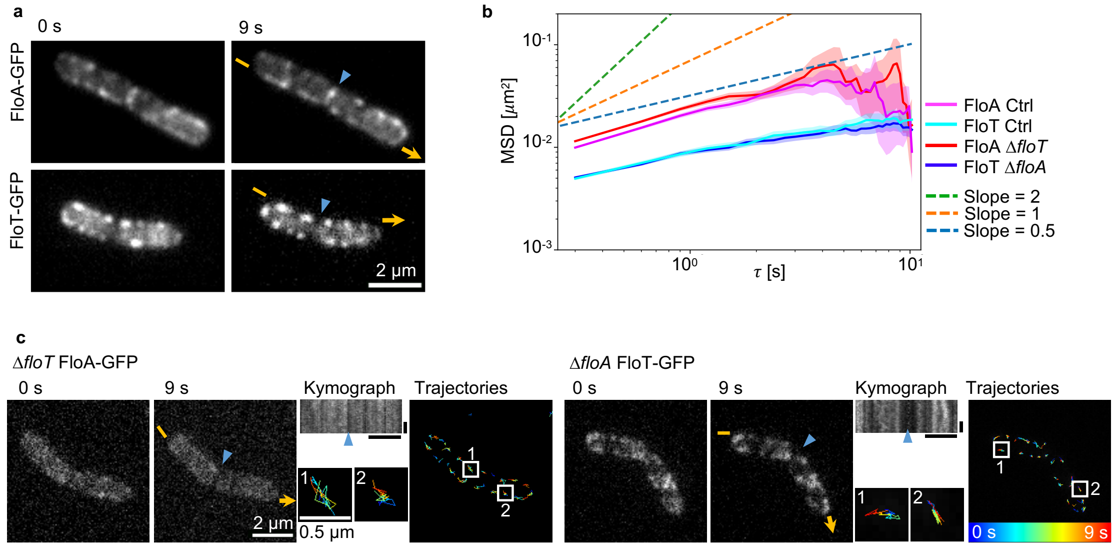


**Supplemental Figure S3: Flotillin mobility is independent of the presence of the other flotillin** (Related to main Figure 1)**. a)** Fluorescence microscopy images showing the subcellular localization of GFP-labeled FloA (top) and FloT (bottom) at 0 s and 9 s. The same cells were used to analyze flotillin dynamics in Fig. 1e and f. The kymographs in Fig. 1e were generated by tracing the membrane signal marked with yellow arrows. Cell poles of neighboring cells are marked with blue triangles. **b)** Plot showing the MSD analysis of FloA in the absence of FloT (Δ*floT*), and FloT in the absence of FloA (Δ*floA*). Plot shows the averages with shaded area representing 95% bootstrap confidence intervals (N≥1538 trajectories). **c)** Time-lapse fluorescence microscopy images showing the movement of FloA-GFP (left panel) and FloT-GFP (right panel) assemblies in the absence of the other flotillin operon, FloA-GFP in Δ*floT* and FloT-GFP in Δ*floA*. Flotillin foci were followed over 9 s (left, first and second column show 0 s and 9 s respectively). The membrane signal marked by yellow arrows was traced to generate kymographs (center column) (horizontal scale bars represent 2 µm, vertical scale bars represent 3 s). The trajectories corresponding to flotillin movements are shown in the right column. Representative trajectories are highlighted. Colors in trajectories indicate elapsed time, from blue=0 s to red=9 s. Cell poles of neighboring cells are marked with blue triangles. The flotillins do not substantially influence each other's movement.

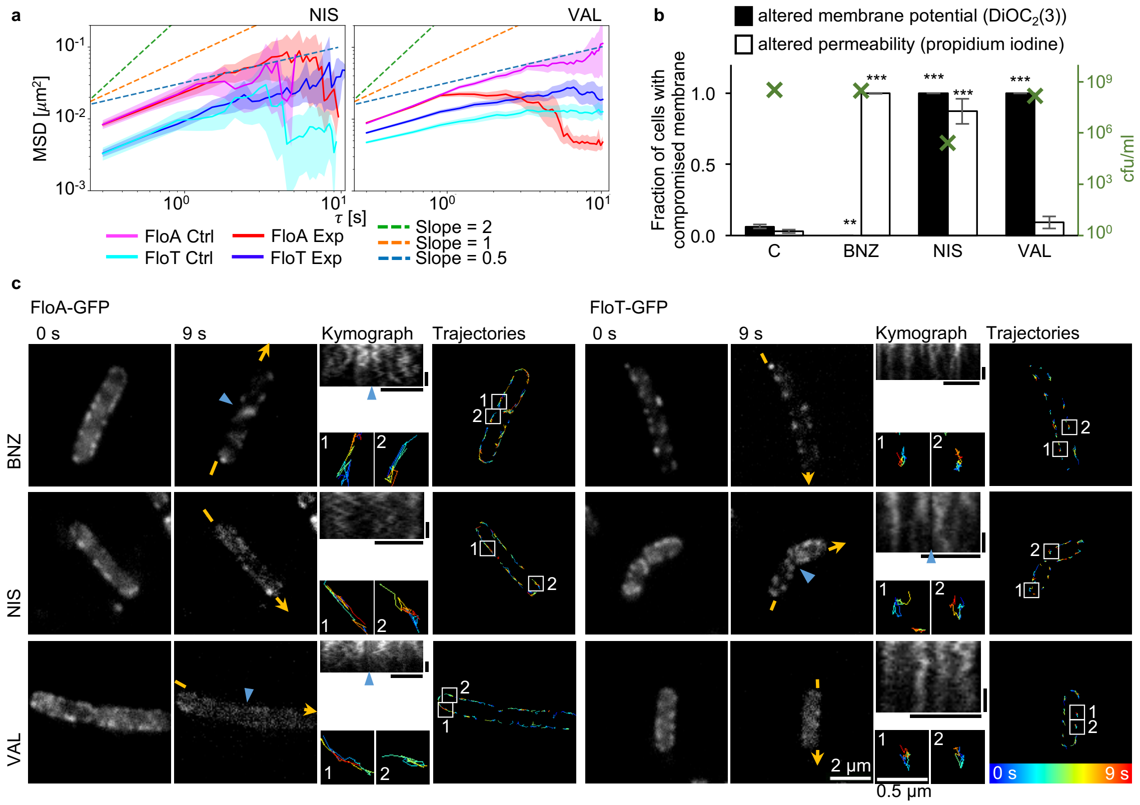


**Supplemental Figure S4: Changes in membrane properties do not alter flotillin mobility** (Related to main Figure 1)**.** *B. subtilis* cells were treated with different membrane-property affecting compounds. **a)** Plots showing the MSD analysis of FloA and FloT assemblies after altering membrane properties using different agents. Nisin (NIS) is a pore former and thus perturbs membrane integrity including transmembrane gradients (XXX) and valinomycin (VAL) abolishes the membrane potential by selectively transporting potassium ions across the membrane (XXX). No differences in flotillin mobility were detected after NIS treatments (left panel, N≥73 trajectories). A detailed analysis is presented in Supp. Fig. S12e. VAL treatment did not influence FloA mobility, but increased FloT mobility (right panel, N≥1351 trajectories). This can be explained by the mislocalization of MreB upon abolishment of the membrane potential, as discussed in detail in Supp. Fig. S18. Nevertheless, NIS also abolished the membrane potential but did not reveal this increase in FloT mobility. Plots show the averages with shaded area representing 95% bootstrap confidence intervals. Ctrl=control condition, Exp=experimental condition (specified on the top). **b)** CFU/ml count (green crosses, referred to right *y*-axis) and cell staining assays (left *y*-axis) to monitor bacterial membrane perturbation. Propidium iodine (red fluorescence signal) was used as a staining probe. It only accumulates in the cytoplasm of cells with compromised membrane permeability. The cytoplasmic dye $DiOC_2(3)$ shifts from green to red fluorescence if membrane integrity and thus membrane potential is not compromised[1]. The fraction of cells with green signal is depicted, indicating differences in the membrane potential. Bar chart shows means±SD (N=250 cells), ** $p<0.01$, *** $p<0.001$. **c)** Time-lapse fluorescence microscopy images showing the movement of FloA-GFP (left panel) and FloT-GFP (right panel) assemblies after treatment. Flotillin foci were followed every 300 ms over 9 s (left, first and second column show 0 s and 9 s respectively). The membrane signal marked by yellow arrows was traced to generate kymographs (center column) (horizontal scale bars represent 2 μm, vertical scale bars represent 3 s). The trajectories corresponding to flotillin movements are shown in the right column. Representative trajectories are highlighted. Colors in trajectories indicate elapsed time, from blue=0 s to red=9 s. Membrane perturbation did not influence flotillin dynamics. C=control untreated cultures; BNZ=benzyl alcohol, NIS=nisin, VAL=valinomycin.

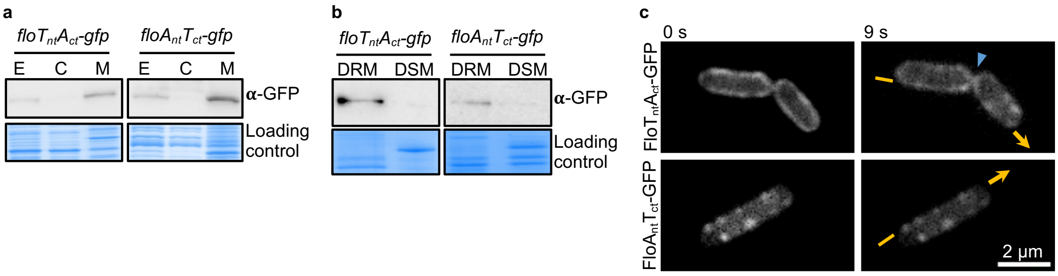


**Supplemental Figure S5: Subcellular localization of chimeric flotillins in the DRM fraction** (Related to main Figure 2)**. a)** Cellular fractionation coupled to the immunodetection of FloT$_{nt}$A$_{ct}$-GFP (left) and FloA$_{nt}$T$_{ct}$-GFP (right) signals in the membrane fraction, performed using α-GFP antibodies. A Coomassie-stained gel is shown as a protein loading control. E=whole cell extract, C=cytosol, M=membrane. **b)** Fractionation of the membrane into DRM and DSM fractions, plus immunodetection of FloT$_{nt}$A$_{ct}$ (left) and FloA$_{nt}$T$_{ct}$ (right), revealed signals mostly in the DRM fraction. **c)** Time-lapse fluorescence microscopy images at 0 s and 9 s, showing the movement of GFP-labeled FloT$_{nt}$A$_{ct}$-expressing (top panel) and FloA$_{nt}$T$_{ct}$-expressing (bottom panel) strains. The kymographs of Figure 2e were generated by tracing the membrane signal marked by yellow arrows. Cell poles of neighboring cells are marked with blue triangles. This experiment revealed the trajectories of FloT$_{nt}$A$_{ct}$ and FloA$_{nt}$T$_{ct}$ as shown in Figure 2f.

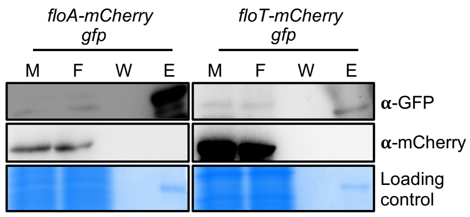


**Supplemental Figure S6: Flotillins coelute with PBP3 and DltD** (Related to main Figure 4)**.** Pulldown control experiments that analyzed the coelution of FloA-mCherry or FloT-mCherry with untagged GFP. A Coomassie-stained gel is shown as a protein loading control. Immunodetection of mCherry in the coeluted samples gave no positive results, confirming that the interaction of flotillins with GFP-PBP3 and GFP-DltD depends on PBP3 and DltD respectively. M=whole membrane fraction, F=flowthrough, W=washed material, E=elution fraction.

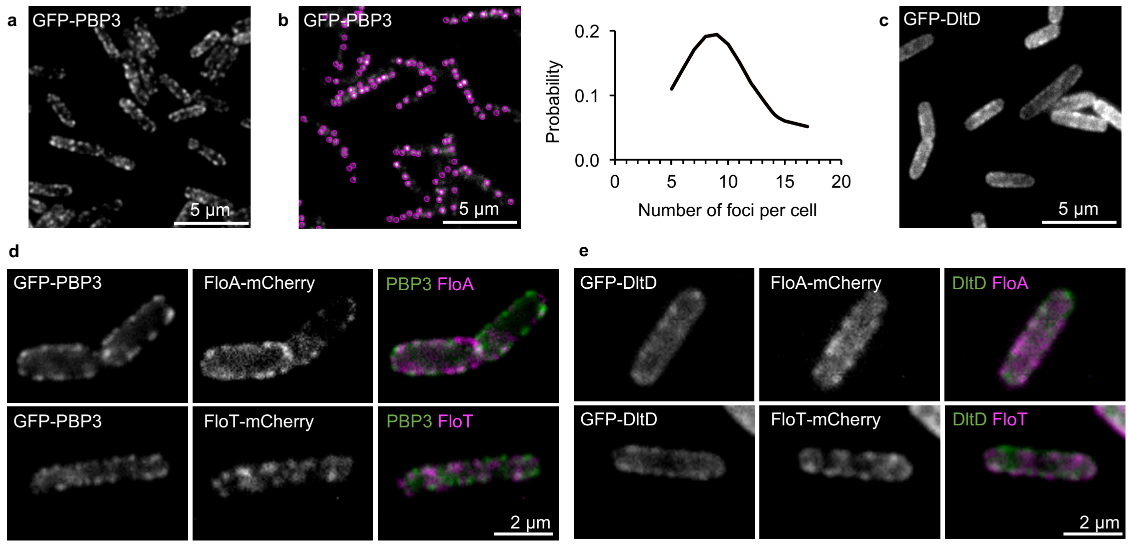


**Supplemental Figure S7: Localization studies of flotillins and their interaction partners PBP3 and DltD** (Related to main Figure 4)**. a)** Fluorescence microscopy images of GFP-PBP3-labeled cells showing a punctate localization pattern for PBP3. **b)** Spot detection exemplified with TIRF images of Fig 4c (left) and quantification of the relative abundance of visible fluorescence foci for PBP3 (N=50 cells) in TIRFM images (right). PBP3 localizes in 9 foci on average. **c)** Fluorescence microscopy images of GFP-DltD-labeled cells showing a heterogeneous localization for DltD. **d)** Fluorescence microscopy experiments to detect the colocalization of PBP3 and flotillins, using GFP-PBP3 + FloA-mCherry (top panel) or GFP-PBP3 + FloT-mCherry (bottom panel) double-labeled strains. The PBP3-FloA colocalization signal was detected more frequently than the PBP3-FloT signal. **e)** Fluorescence microscopy experiments to detect the colocalization of DltD and flotillins, using GFP-DltD + FloA-mCherry (top panel) and GFP-DltD + FloT-mCherry (bottom panel) double-labeled strains. The DltD-FloT colocalization signal was detected more frequently than the DltD-FloA signal.

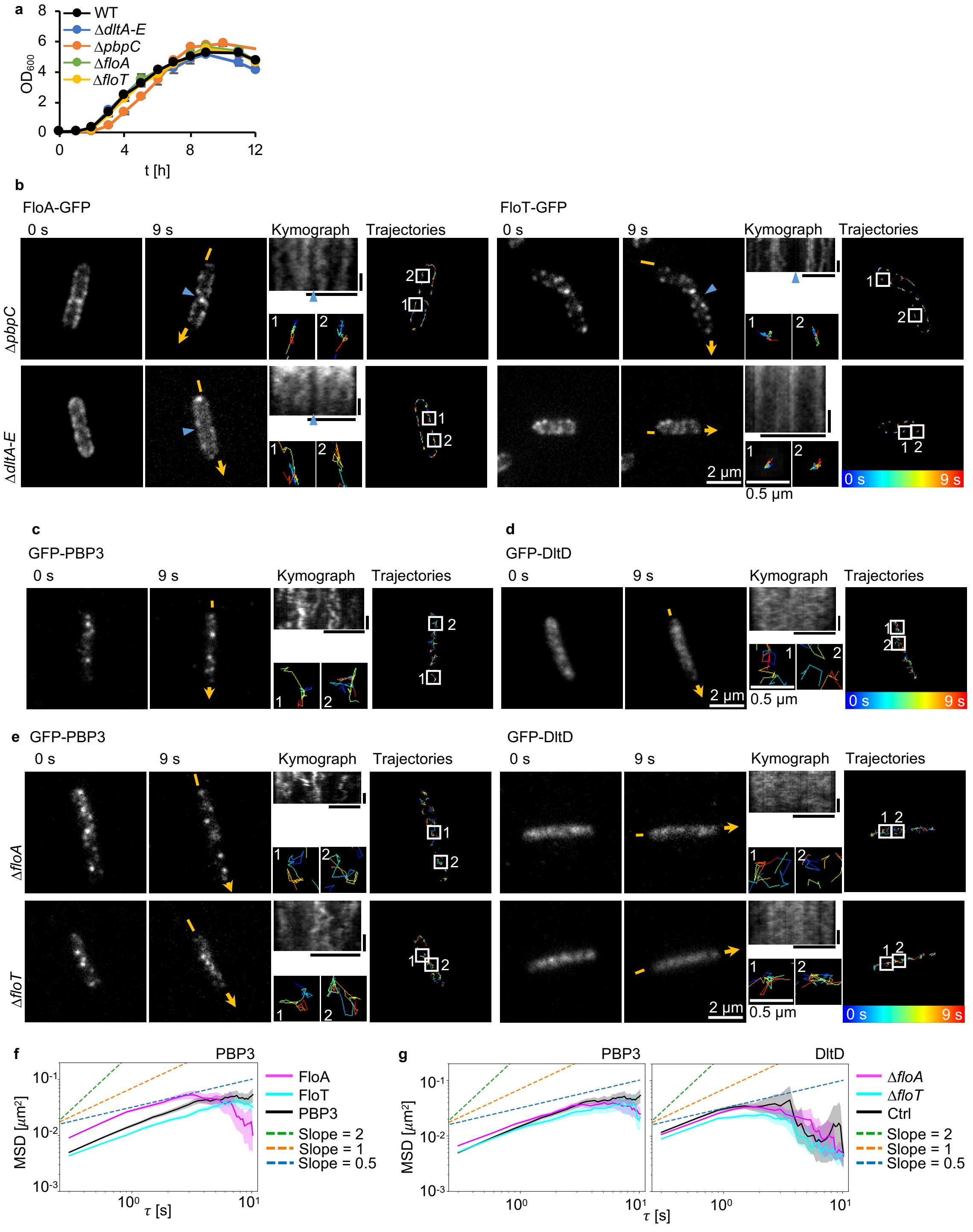


**Supplemental Figure S8: Mutants affecting the mobility of the flotillins and their interaction partners** (Related to main Figure 4)**. a)** Growth curves for WT, Δ*pbpC*, Δ*dltA-E*, Δ*floA* and Δ*floT* strains in LB medium show no differences. **b)** Fluorescence microscopy analyses of FloA (left panel) and FloT (right panel) dynamics in Δ*pbpC* and Δ*dltA-E* backgrounds. Images were acquired every 300 ms for 9 s;

localization patterns at 0 s and 9 s (left, first and second columns) and kymographs (center column) (horizontal scale bars represent 2 μm, vertical scale bars represent 3 s), as well as the trajectories of the individual flotillin foci (right column), are shown. Representative trajectories are highlighted in more detail. Colors indicate elapsed time from blue=0 s to red=9 s. Cell poles are marked with blue triangles. The mobility of FloT was substantially reduced in the Δ*dltA-E* background. **c)** TIRF microscopy images monitoring PBP3 movement; images were captured every 300 ms over 9 s. The localization patterns at 0 s and 9 s (left, first and second column), kymographs for the PBP3 signal along the longitudinal axis of the cell marked by yellow arrows (center column) (horizontal scale bars represent 2 μm, vertical scale bars represent 3 s) and the trajectories of individual PBP3 foci (right column) are shown. According to the kymographs the mobility of PBP3 revealed tracks showing lateral displacements. Trajectories cover larger membrane areas with occasional spatial restrictions. PBP mobility is comparable to that of FloA and FloT. **d)** TIRF microscopy images showing the heterogeneous localization of DltD in membrane regions. The detection of foci using Fiji software was not as efficient as when punctate localization patterns were analyzed. To analyze the movement of DltD, images were captured every 300 ms over 9 s. The localization patterns at 0 s and 9 s (left, first and second column) are shown. Kymographs were generated by tracing the membrane signal marked by yellow arrows (center column) and the trajectories of individual DltD foci (right column) are shown. According to the kymographs, and given the short trajectories, the DltD signal reveals rapid movement, although the differences in the type of the signal precluded any comparison with signals distributed in discrete membrane foci. **e)** TIRF microscopy analyses of PBP3 (left panel) and DltD (right panel) movement in Δ*floA* and Δ*floT* backgrounds. Localization pattern, kymographs and trajectories were analyzed as described above. PBP3 mobility increased in the Δ*floA* background and DltD mobility decreased in the Δ*floT* background. **f)** Plot showing the MSD analysis of FloA, FloT and PBP3. Plot shows the averages with shaded area representing 95% bootstrap confidence intervals (N≥1275 trajectories). PBP3 shows diffusive behavior, and the PBP3 diffusion coefficient settles between FloA and FloT, slower than FloA and faster than FloT. **g)** Plots showing the MSD analysis of PBP3 (left panel, N≥1075 trajectories) and DltD (right panel, N≥1259 trajectories) in Δ*floA* and Δ*floT* mutants. A minor increase of PBP3 in Δ*floA* and a minor decrease of DltD in Δ*floT* is visible. Ctrl=control condition (specified on the top). Plots show the averages with shaded area representing 95% bootstrap confidence intervals.

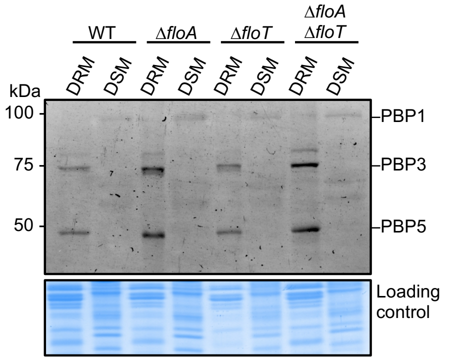


**Supplemental Figure S9: PBP3 protein levels increased in the Δ*floA* mutant** (Related to main Figure 4)**.** DRM/DSM fractionation of Bocillin-FL-stained membrane fractions to detect PBP localization and abundance in WT, Δ*floA*, Δ*floT* and Δ*floA* Δ*floT* backgrounds. A Coomassie-stained gel is shown as a protein loading control. The levels of PBP3 and PBP5 increased in strains lacking FloA.

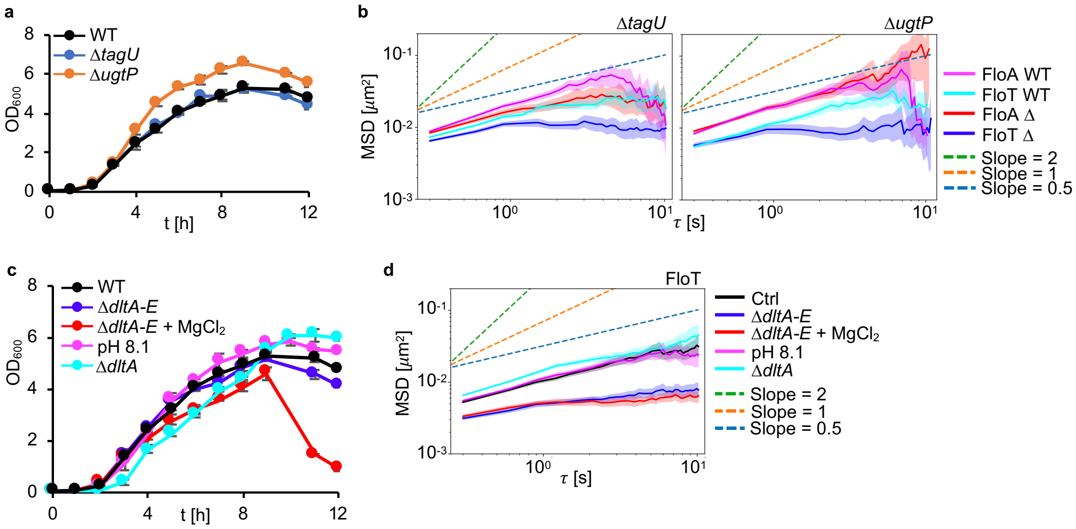


**Supplemental Figure S10: The reduced mobility of FloT in the Δ*dltA-E* mutant depends on DltD** (Related to main Figure 4)**. a)** DltA-E modify anionic polymers (wall teichoic acids [WTA] and lipoteichoic acids [LTA]) of the cell wall with D-alanyl residues. To understand if differences in FloT mobility upon inhibition of DltA-E, result from differences in anionic polymers, we used Δ*tagU* and Δ*ugtP* mutations. These are approximations for inhibition of WTA and LTA synthesis, respectively, as complete mutations result in severe growth defects. Growth curves for WT, Δ*tagU* and Δ*ugtP* strains in LB medium show similar growth. **b)** Plots showing the MSD analysis of FloA and FloT in Δ*tagU* (left panel) and Δ*ugtP* (right panel) strain backgrounds (N≥488 trajectories). No differences in flotillin mobility can be observed after inhibition of WTA or LTA synthesis. **c)** As Δ*tagU* and Δ*ugtP* are only approximations, we used different approaches to confirm, that the reduced FloT mobility does not result from the differences in cell wall charges but from the absence of DltD. FloT interacts with DltD and the diffusion coefficient of FloT is reduced in the Δ*dltA-E* mutant. To confirm that the reduced mobility in the mutant results from the absence of DltD, analyses were performed with the following conditions: i) addition of 25 mM $MgCl_2$ to the culture to counteract the absence of positively charged D-alanyl residues in the cell wall, ii) growth at pH 8.1, which leads to D-alanyl residue loss from the cell wall[6], iii) use of the Δ*dltA* mutant, which does not incorporate D-alanyl residues into the cell wall[7]. Similar growth rates were detected under all conditions. **d)** Plot showing the MSD analysis of FloT in conditions i to iii. The reduced mobility of FloT in the Δ*dltA-E* background results from the absence of DltD and not from cell wall differences. Plots show the averages with shaded area representing 95% bootstrap confidence intervals (N≥1134 trajectories).

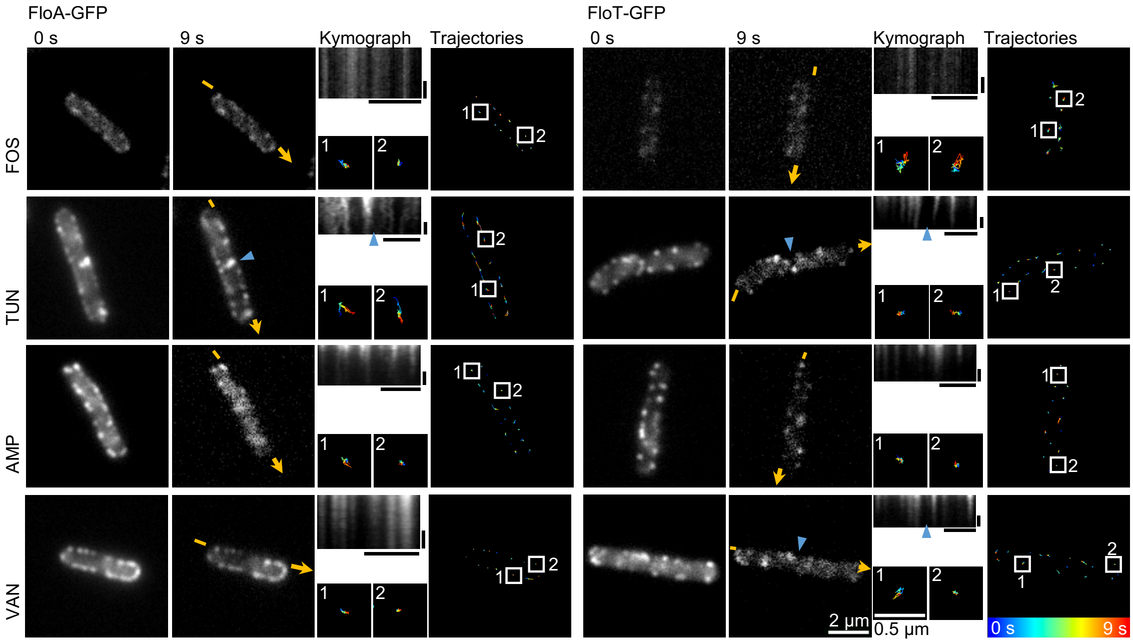


**Supplemental Figure S11: The dynamic behavior of FloA and FloT is reduced upon the inhibition of cell wall synthesis** (Related to main Figure 4)**.** Fluorescence microscopy analyses of FloA (left panel) and FloT (right panel) movement after the inhibition of cell wall synthesis. Images were acquired every 300 ms for 9 s. The localization patterns at 0 s and 9 s (left, first and second columns) and kymographs (center column) (horizontal scale bars represent 2 µm, vertical scale bars represent 3 s), as well as the trajectories of the individual flotillin foci (right column), are shown. Representative trajectories are highlighted in more detail. Colors indicate elapsed time from blue=0 s to red=9 s. Cell poles of neighboring cells are marked with blue triangles. FOS=fosfomycin, TUN=tunicamycin, AMP=ampicillin, VAN=vancomycin.

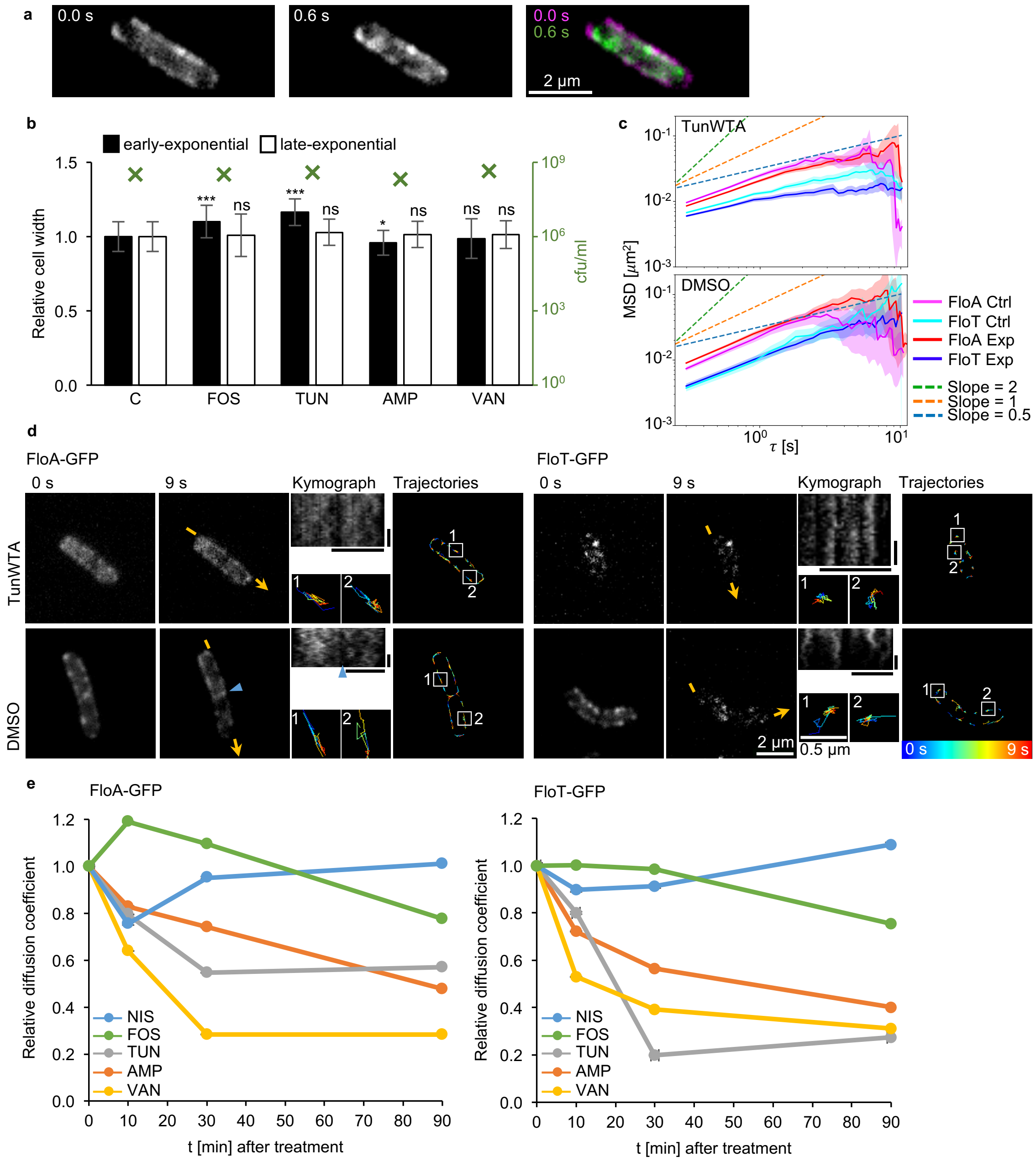


**Supplemental Figure S12: Flotillin movement is reduced due to the inhibition of cell wall synthesis** (Related to main Figures 2 and 4). **a)** Cell death causes plasmolysis[8], visible as cell shrinkage in these fluorescence microscopy images showing a cell before and after plasmolysis. **b)** Cell width was recorded upon the inhibition of cell wall synthesis (N≥28 cells) (left *y*-axis). The CFU/ml count was also monitored (green crosses, referred to right *y*-axis). No reduction in cell counts were detected. Cell widths are not reduced either, except for a small decrease after AMP treatments. These results confirm that differences in flotillin mobility are due to the inhibition of cell wall synthesis. Note that an increase in cell width has been reported in cells defective for wall teichoic acid (WTA) synthesis[9]. Bar chart shows means±SD. **c)** Plots showing the MSD analysis of FloA and FloT in control experiments. Control experiments using the solvent (DMSO 2%), or using TUN at a reduced concentration (TunWTA, 0.025 μg/ml) were performed. TUN inhibits the enzymatic activity of MraY and TagO to different degrees. TagO, which is involved in WTA synthesis, is inhibited at low concentrations of TUN whereas inhibition of MraY, which is involved in lipid II synthesis, mainly occurs at higher concentrations[10–12]. Compared to the effect of TUN treatment, only minor differences in the diffusion coefficients of FloA and FloT were observed after TunWTA (top, N≥1225 trajectories) or DMSO (bottom, N≥151 trajectories) treatments. This confirms that the differences in flotillin mobility after TUN treatment are due to the inhibition of cell wall synthesis. Plots show the averages with shaded area representing 95% bootstrap confidence intervals. Ctrl=control condition, Exp=experimental condition (specified in the top left corner of each plot). **d)** Fluorescence microscopy analysis of FloA (left panel) and FloT (right panel) dynamics after treatments with TunWTA and DMSO. Images were acquired every 300 ms for 9 s; localization patterns at 0 s and 9 s (left, first and second columns) and kymographs (center column) (horizontal scale bars represent 2 μm, vertical scale bars represent 3 s), as well as the trajectories of the individual flotillin foci (right column), are shown. Representative trajectories are highlighted in more detail. Colors indicate elapsed time from blue=0 s to red=9 s. Cell poles of neighboring cells are marked with blue triangles. **e)** NIS forms pores in the membrane by binding to and sequestering the cell wall synthesis precursor lipid II[13]. Nevertheless, no differences in flotillin mobilities were observed after NIS treatment (Supp. Fig. S4). It may be that upon NIS treatment, lipid II is sequestered and cell wall synthesis is temporarily inhibited until *de novo* synthesized lipid II becomes available. FloA (left) and FloT (right) movements were thus monitored at different time points (10 min, 30 min and 90 min) after inhibition with NIS (N≥73 trajectories), FOS (N≥273 trajectories), TUN (N≥83 trajectories), AMP (N≥84 trajectories) or VAN (N≥208 trajectories). A reduction in flotillin mobility upon NIS treatment was seen only at early time points. In contrast, flotillin mobility gradually decreased with incubation time in cells with inhibited cell wall synthesis. Graph shows relative diffusion coefficient. Diffusion coefficient $D = MSD(\tau = 0.3) / (4 * 0.3\ s)$ was set to 1 for each of the 0 minute conditions. Means±sem. C=control untreated cultures, NIS=nisin, FOS=fosfomycin, TUN=tunicamycin, AMP=ampicillin, VAN=vancomycin. ns=not significant, * $p<0.05$, *** $p<0.001$.

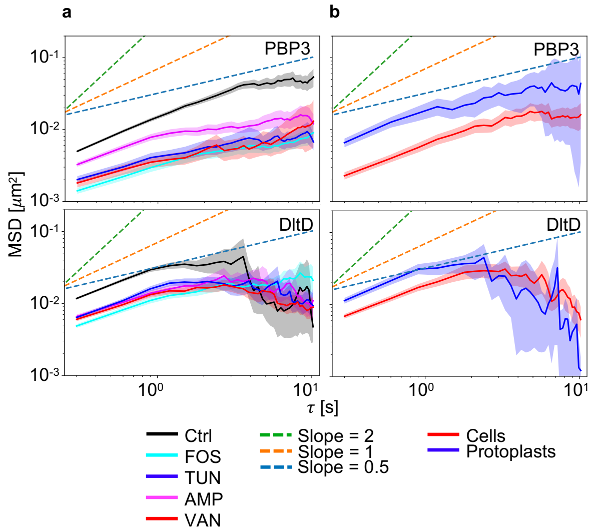


**Supplemental Figure S13: The mobility of PBP3 and DltD is reduced upon cell wall synthesis inhibition and increased in protoplasts** (Related to main Figure 4)**.** Plots showing MSD analysis of PBP3 (top panels) and DltD (bottom panels) in **a)** cells treated with different cell-wall synthesis inhibitors or in **b)** protoplasts. Similar to FloA and FloT, PBP3 and DltD mobility is reduced in cell wall synthesis inhibitory conditions (N≥538 trajectories) and increased in protoplasts (N≥102 trajectories). FOS=fosfomycin, TUN=tunicamycin, AMP=ampicillin, VAN=vancomycin. Plots show the averages with shaded area representing 95% bootstrap confidence intervals.

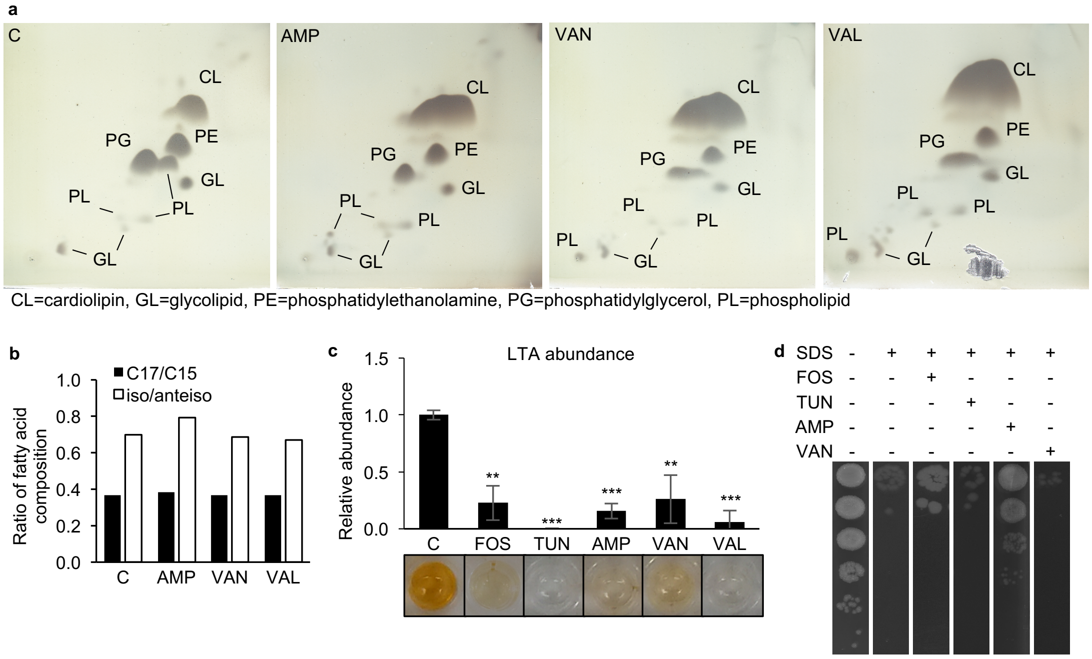


**Supplemental Figure S14: Membrane properties do not account for differences in flotillin mobility** (Related to main Figure 4)**.** Analysis of membrane composition and properties after treating cells with antibiotics that affect flotillin mobility (FOS, TUN, AMP, VAN) in relation to compounds that do *not* affect flotillin mobility (VAL). **a)** 2D-TLC analysis of membrane polar lipids showed an increase in CL levels with all treatments independent of variations in flotillin mobility. CL=cardiolipin, GL=glycolipid, PE=phosphatidylethanolamine, PG=phosphatidylglycerol, PL=phospholipid. **b)** Gas chromatography coupled to mass spectrometry analyses of membrane fatty acids showed no alterations in the ratio of C17/C12 or iso/anteiso that correlated with differences in flotillin mobility. **c)** ELISA assays to monitor the abundance of lipoteichoic acid (LTA) in the membranes of treated cells. No correlation was seen between LTA levels and flotillin mobility. Bar chart shows means±SD (N=3), * $p<0.1$, ** $p<0.01$, *** $p<0.001$. **d)** Cell survival after SDS (0.05%) treatment of cells pretreated with antibiotics showed no differences in relation to flotillin mobility changes. FOS=fosfomycin, TUN=tunicamycin, AMP=ampicillin, VAN=vancomycin, VAL=valinomycin

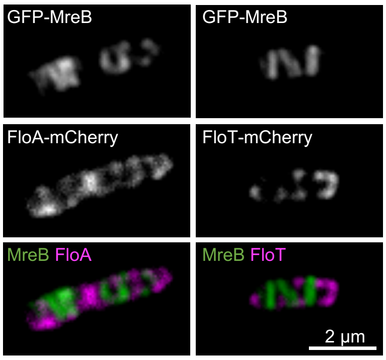


**Supplemental Figure S15: FloA and FloT do not colocalize with MreB** (Related to main Figure 5)**.** Fluorescence microscopy experiments showing colocalization of the MreB signal with FloA or FloT signals using the GFP-MreB + FloA-mCherry (left) or GFP-MreB + FloT-mCherry (right) double-labeled strains. The flotillin and MreB signals exclude one another.

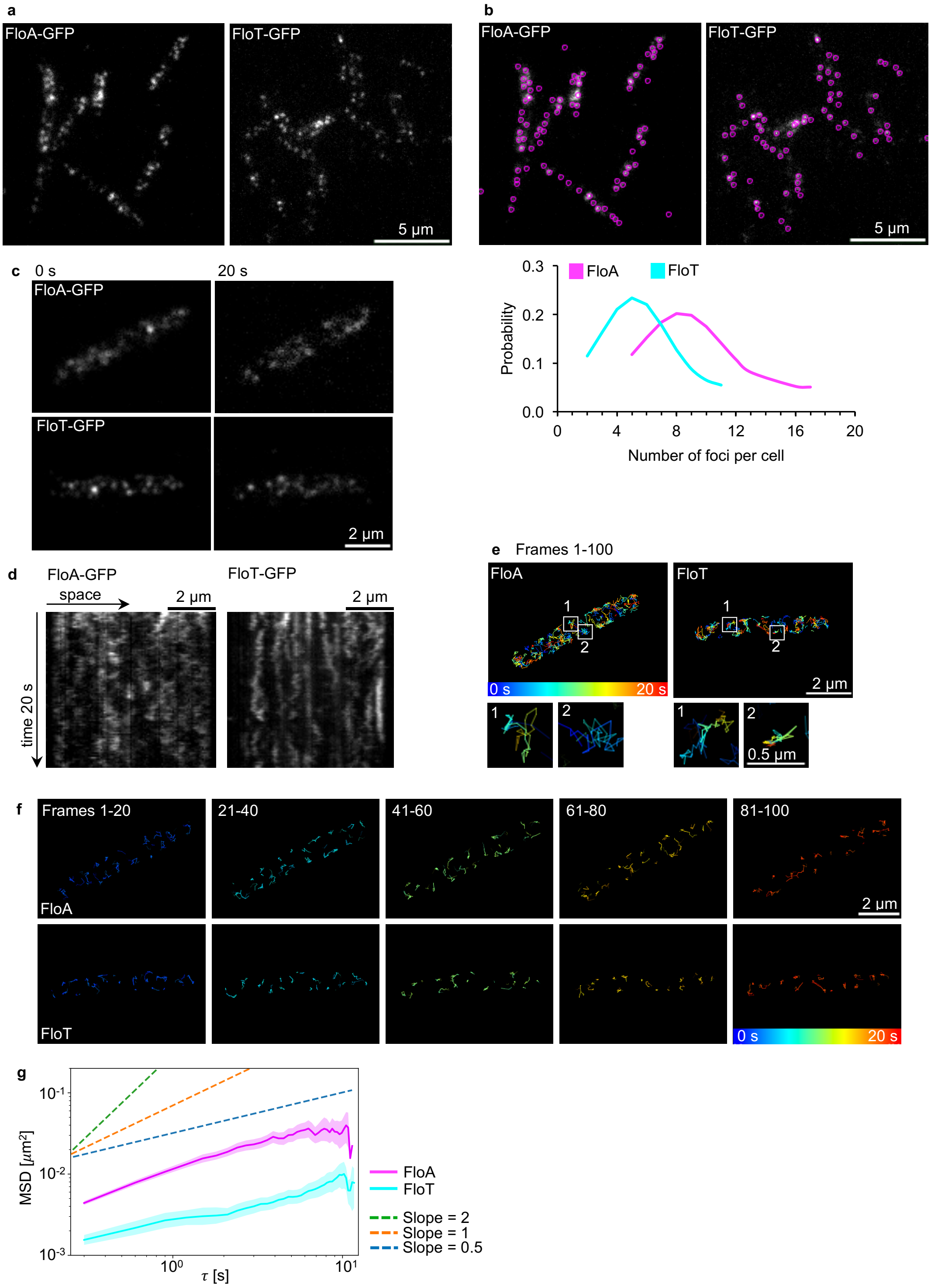


**Supplemental Figure S16: Localization and movement of FloA and FloT using TIRF microscopy** (Related to main Figure 5). **a)** TIRF microscopy images showing the subcellular localization pattern of GFP-labeled FloA (left panel) and FloT (right panel) foci. **b)** Spot detection exemplified with images of a) (top) and quantification of the relative abundance of fluorescence foci for FloA and FloT assemblies (bottom). On average, FloA localized in 8 visible foci and FloT in 5 visible foci (N=50 cells). **c)** TIRF microscopy analysis of FloA (top panel) and FloT (bottom panel) movements. Images were captured every 200 ms over 20 s to determine mobility. The localization patterns at 0 s and 20 s are shown. **d)** Kymographs of the membrane signal along the longitudinal cell axis of FloA-GFP (left) and FloT-GFP (right) are shown. **e)** The trajectories the individual flotillin foci followed over 20 s are shown. FloA showed spatially restricted movement as well as movement over large membrane areas. FloT mainly showed spatially restricted movement that concentrated in smaller membrane areas. FloA covers more overall membrane area than FloT within these 20 s. Representative trajectories are highlighted in more detail. Colors indicate elapsed time from blue=0 s to red=20 s. **f)** Mobility of FloA-GFP (top) and FloT-GFP (bottom) represented in local trajectories flotillin foci follow. Numbers indicate different frame ranges. **g)** Plot showing the MSD analysis of FloA and FloT detect the same tendency as similar to that obtained by fluorescence microscopy, albeit at lower diffusion coefficients. FloA diffuses faster than FloT. Plot shows the averages with shaded area representing 95% bootstrap confidence intervals (N≥765 trajectories).

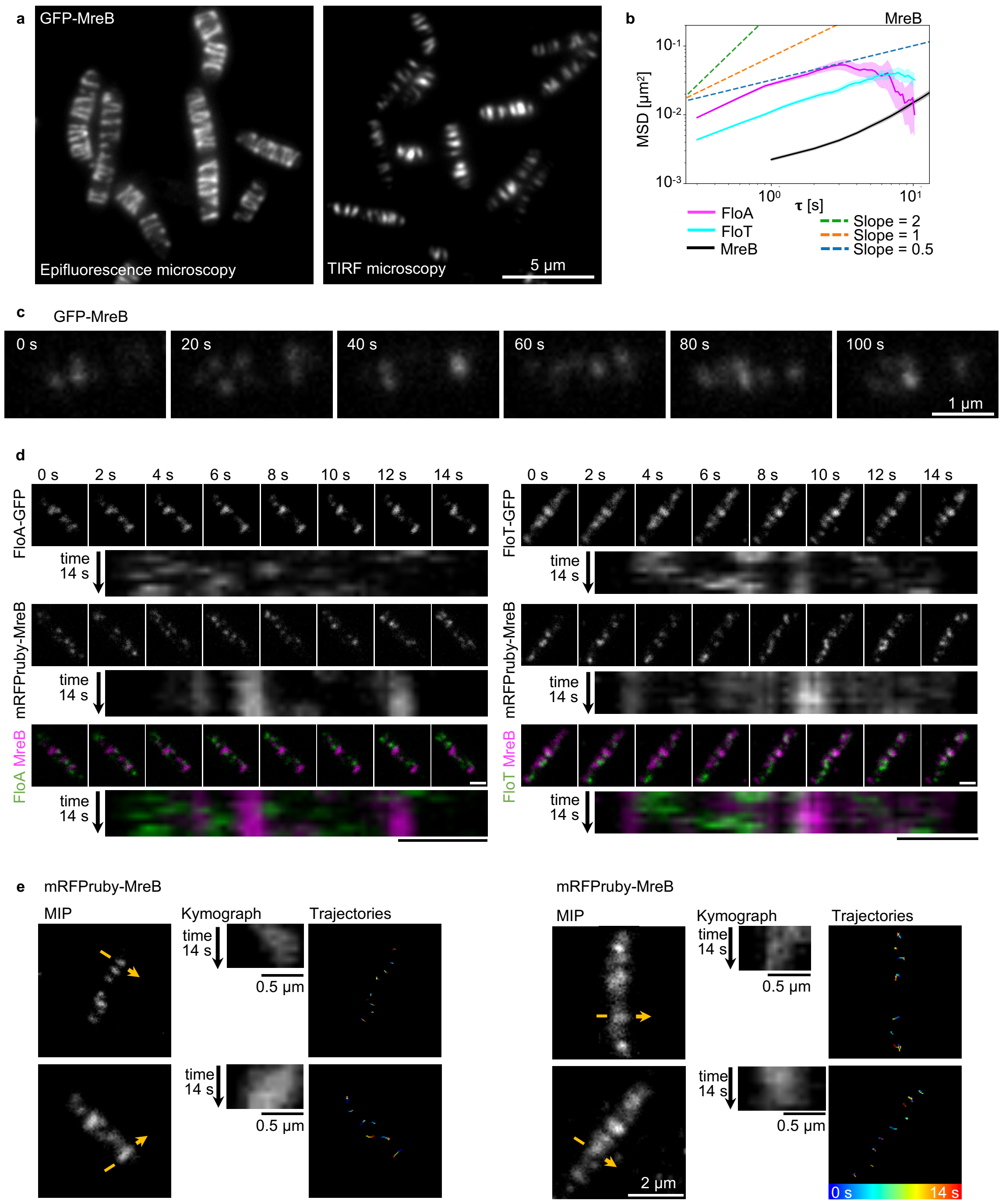


**Supplemental Figure S17: Localization and mobility of MreB using TIRF microscopy.** (Related to main Figure 5) **a)** Subcellular localization pattern of GFP-MreB using epifluorescence (left panel) or TIRF microscopy (right panel). **b)** Plot showing the MSD analysis of GFP-labeled FloA, FloT and MreB cells. MreB moves more slowly than the flotillins. Plot shows the averages with shaded area representing 95% bootstrap confidence intervals (N≥850 trajectories). **c)** The mobility of MreB can only be monitored using TIRF microscopy; results are shown as sequences of 20 s time intervals for the same cell as discussed in Fig. 5e. **d)** Additional example of time sequences of FloA-GFP (left), FloT-GFP (right) and mRFPruby-MreB double-labeled strains. Images were acquired with simultaneous TIRF microscopy (images captured every 2 s over 14 s) and kymographs corresponding to the signal of the longitudinal axis of the cell are shown. Scale bars represent 1 µm. **e)** TIRF microscopy equipped for simultaneous image acquisition showing MreB mobility in experiments of Fig. 5f and Supp. Fig. S17d. Confirmation of MreB mobility is presented for Fig. 5f (here top, left and right correspond to the respective cells in Fig. 5f) and Fig. S17d (here bottom, left and right correspond to the respective cells in Fig. S17d). Images were taken at 2 s intervals over 14 s; this allowed the MreB movements to be followed but was too large a time scale to capture flotillin movements properly. MIP of MreB (left column), kymographs showing signals perpendicular to the cell axis, as indicated by yellow arrows (center column) and trajectories of individual foci (right column) confirmed habitual MreB mobility. Colors indicate elapsing time, from blue=0 s to red=14 s.

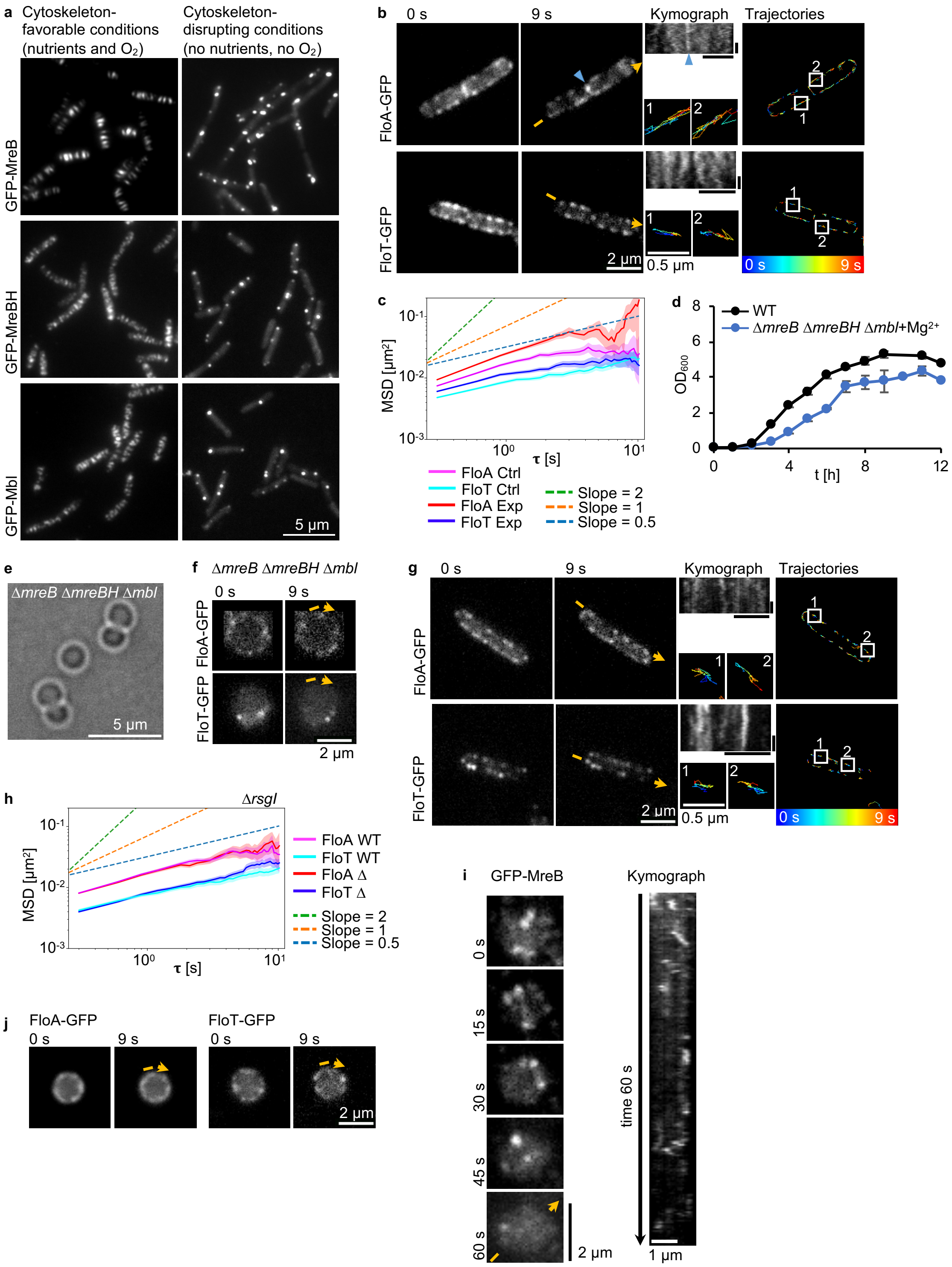


**Supplemental Figure S18: Disorganization or absence of actin-homolog filaments increased flotillin movement** (Related to main Figure 6)**. a)** *B. subtilis* cells contain the actin-homolog proteins MreB, MreBH and Mbl. Their filaments are disrupted upon disturbance of the membrane potential[14] by limiting the access to nutrients and oxygen (right). For correct MreB localization, access to nutrients and oxygen must be maintained (left). **b)** Fluorescence microscopy analysis of FloA-GFP (top) and FloT-GFP (bottom) movement under cytoskeleton-disrupting conditions (blockage of nutrient and oxygen availability). Images were acquired every 300 ms for 9 s; localization patterns at 0 s and 9 s (left, first and second columns) and kymographs (center column) (horizontal scale bars represent 2 µm, vertical scale bars represent 3 s), as well as the trajectories of the individual flotillin foci (right column), are shown. Representative trajectories are highlighted in more detail. Colors indicate elapsed time from blue=0 s to red=9 s. Cell poles of neighboring cells are marked with blue triangles. **c)** Plot showing the MSD analysis of FloA and FloT in cytoskeleton-disrupting conditions. Plot shows the averages with shaded area representing 95% bootstrap confidence intervals (N≥1785 trajectories). Ctrl=control condition (cytoskeleton-favorable), Exp=experimental condition (cytoskeleton-disrupting). The diffusion coefficients of FloA and FloT increase in conditions where the actin-homolog cytoskeleton is disrupted. **d)** Mutants lacking all three actin-homologs are only viable with an additional mutation in *rsgI*. This strain does not grow in LB medium unless $MgSO_4$ is added. **e)** Bright-field microscopy images of this strain show that it grows as spheres. **f)** Movement of FloA-GFP (top) and FloT-GFP (bottom) in actin-homolog mutant strains was analyzed by fluorescence microscopy. Images were acquired every 300 ms for 9 s and localization patterns at 0 s and 9 s monitored. The kymographs in Fig. 6a were generated by tracing the membrane signal marked with yellow arrows. **g)** Movement of FloA-GFP (top) and FloT-GFP (bottom) in Δ*rsgI* mutant control strains was analyzed by fluorescence microscopy. Images were acquired every 300 ms for 9 s; localization patterns at 0 s and 9 s (left, first and second columns) and kymographs (center column) (horizontal scale bars represent 2 µm, vertical scale bars represent 3 s), as well as the trajectories of the individual flotillin foci (right column), are shown. Representative trajectories are highlighted in more detail. Colors indicate elapsing time from blue=0 s to red=9 s. **h)** Plot showing the MSD analysis of FloA and FloT in Δ*rsgI* strain background. No differences in flotillin mobility can be observed. Plot shows the averages with shaded area representing 95% bootstrap confidence intervals (N≥1196 trajectories). **i)** The cell wall of GFP-MreB labeled strains was digested in osmotically stabilizing conditions to generate protoplasts. The mobility of MreB was monitored with TIRF microscopy every 300 ms over 60 s and is presented in Fig. 6c, d. Results are shown as sequences of 15 s time intervals (left) and kymograph analysis (right). The mobility of MreB in protoplasts is disoriented and increased. **j)** Time lapse fluorescence microscopy analysis of FloA (top) and FloT (bottom) movement in protoplasts. Images were acquired every 300 ms for 9 s; localization patterns at 0 s and 9 s are shown. The kymographs in Fig. 6e were generated by tracing the membrane signal that is marked with yellow arrows.

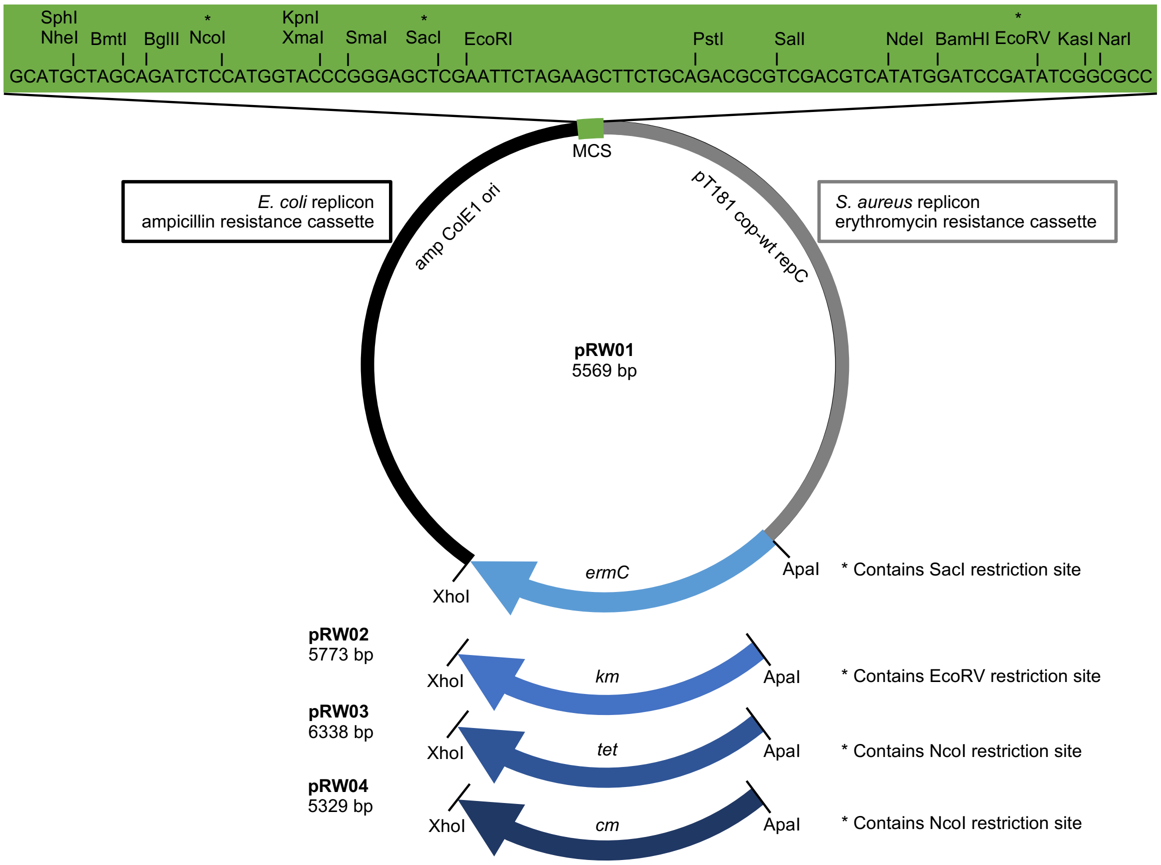


**Supplemental Figure S19: Design of *B. subtilis* replicative plasmids** (See Materials and Methods)**.** The shuttle vector pJL-sar-gfp (ampicillin resistance for proliferation in *E. coli* and erythromycin resistance for selection in Gram-positive bacteria)[15] was used as a backbone to replace the *sar-gfp* insert with a multiple cloning site. Several versions of the plasmid with different antibiotic resistance cassettes for Gram-positive bacteria were then constructed.

### Supplemental Tables

Supplemental Table S1: Activity, MIC and concentration of antibiotics used in this work.

| | Target | Compound[a] | MIC[b] | Concentration[c] | Mechanism of action | Reference |
|---|---|---|---|---|---|---|
| Membrane | Lipid bilayer | Benzyl alcohol (BNZ) | nd | 30 mM | Increases membrane fluidity. | [16] |
| | Lipid II | Nisin (NIS) | nd | 30 µM | Lantibiotic causing pore formation by interaction and sequestration of Lipid II. | [17] |
| | $K^+$-ions | Valinomycin (VAL) | nd | 60 µM[d] | Ionophore selectively transporting potassium ions across the membrane abolishing electrochemical gradients. | [17] |
| Cell wall synthesis | MurA | Fosfomycin (FOS) | 250 µg/ml | 2.5 mg/ml | Enzymatic inhibition of the first step in cell wall synthesis by alkylation of this phosphoenolpyruvate analog with the active site cysteine. | |
| | TagO MraY | Tunicamycin (TUN) | 0.5 µg/ml | 0.025 µg/ml or 2.5 µg/ml | Nucleoside antibiotic inhibiting the glycosylation of UPP with WTA and Lipid II precursors. | [10,11] |
| | Class b PBPs | Ampicillin (AMP) | 100 µg/ml | 1 mg/ml | Beta-lactam penicillin causing irreversible inactivation of class b PBPs by binding to their active site of transpeptidation reaction. | |
| | D-Ala-D-Ala of pentapeptide | Vancomycin (VAN) | 0.5 µg/ml | 5 µg/ml | None-ribosomal peptide sterically inhibiting transpeptidation by binding to D-Ala-D-Ala of the cell wall acceptor pentapeptide. | |

[a] Parenthesis define abbreviations used in this study.
[b] nd = not determined
[c] If available, the same concentrations as in appropriate published studies were used, it not, 10 x MIC was used
[d] Valinomycin was used in the presence of 300 mM KCl and 50 mM Hepes pH 7.5.

Supplemental Table S2: Raw and analyzed global pulldown data (see separate file)

Supplemental Table S3: List of strains used in this study.

| Strain | | Genotype[a] | Reference | Construction |
|---|---|---|---|---|
| ***Escherichia coli*** | | | | |
| DL95 | DH5α | Wild type | [18] | |
| RW65 | DH5α | pDR183 (*amp*/*mls*) | DZ Rudner Lab (HMS, USA) | |
| RW73 | DH5α | pDR183-$P_{yqeZ}$-*floA-gfp* | This study | EcoRI/SphI |
| RW114 | DH5α | pDR183-$P_{yuaF}$-*floT-gfp* | This study | SalI/SphI |
| JS197 | DH5α | pDR183-$P_{yqeZ}$-*floA-mCherry* | [19] | |
| JS182 | DH5α | pDR183-$P_{yuaF}$-*floT-mCherry* | [19] | |
| RW430 | DH5α | pSG1729 (*amp*/*spc*) | [20] | |
| RW494 | DH5α | pSG1729-$P_{xyl}$-*gfp-dltD* | This study | Xho/EcoRI |
| RW189 | DH5α | pJL-sar-gfp (*amp*/*ermC*) | [15] | |
| RW197 | DH5α | pRW01 (*amp*/*ermC*) | This study | SphI/NarI |
| RW420 | DH5α | pRW01-$P_{yqeZ}$-*floA-mCherry* | This study | EcoRI/SphI |
| RW421 | DH5α | pRW01-$P_{yuaF}$-*floT-mCherry* | This study | KpnI/SphI |
| RW199 | DH5α | pRW02 (*amp*/*km*) | This study | XhoI/ApaI |
| RW200 | DH5α | pRW03 (*amp*/*tet*) | This study | XhoI/ApaI |
| RW201 | DH5α | pRW04 (*amp*/*cm*) | This study | XhoI/ApaI |
| RW394 | DH5α | pRW04-$P_{yqeZ}$-*floA-gfp* | This study | EcoRI/SalI |
| RW390 | DH5α | pRW04-$P_{yuaF}$-*floT-gfp* | This study | KpnI/SalI |
| RW391 | DH5α | pRW04-$P_{yuaF}$-*flo*$T_{Nt}A_{Ct}$-*gfp* | This study | KpnI/SalI |
| RW395 | DH5α | pRW04-$P_{yqeZ}$- *flo*$A_{Nt}T_{Ct}$-*gfp* | This study | EcoRI/SalI |
| ***Bacillus subtilis*** | | | | |
| RW3 | PY79 | Wild type | [21] | |
| RW88 | PY79 | Δ*lacA*::$P_{yuaF}$-*floT-gfp* (*mls*) | This study | RW3+RW114 |
| RW77 | PY79 | Δ*lacA*::$P_{yqeZ}$-*floA-gfp* (*mls*) | This study | RW3+RW73 |
| RW45 | PY79 | Δ*lacA*::$P_{yqeZ}$-*floA-mCherry* (*mls*) | [19] | |
| RW48 | PY79 | Δ*lacA*::$P_{yuAF}$-*floT-mCherry* (*mls*) | [19] | |
| RW578 | PY79 | Δ*floA*::*floA-gfp-tet* | This study | Primers A |
| RW579 | PY79 | Δ*floT*::*floT-gfp-tet* | This study | Primers B |
| RW323 | PY79 | Δ*amyE*::$P_{yuaF}$-*flo*$T_{Nt}A_{Ct}$-*gfp* (*spc*) | [19] | |
| RW375 | PY79 | Δ*lacA*::$P_{yqeZ}$-*flo*$A_{Nt}T_{Ct}$-*gfp* (*mls*) | [19] | |
| RW329 | PY79 | Δ*floA*::*cm* | This study | Primers C |
| DL1237 | 168 | Δ*floA*::*spc* | [22] | |
| RW330 | PY79 | Δ*floT*::*tet* | This study | Primers D |
| RW334 | PY79 | Δ*floA*::*cm* Δ*floT*::*tet* | This study | RW329+RW330 |
| RW28 | PY79 | Δ*floA::mls* Δ*floT::spc* | [23] | |
| RW521 | PY79 | Δ*floT*::*tet* Δ*lacA*::$P_{yqeZ}$-*floA-gfp* (*mls*) | This study | RW330+RW73 |
| RW522 | PY79 | Δ*floA*::*cm* Δ*lacA*::$P_{yuaF}$-*floT-gfp* (*mls*) | This study | RW329+RW114 |

| Strain | | Genotype[a] | Reference | Construction |
|---|---|---|---|---|
| RW404 | PY79 | Δ*floA::mls* Δ*floT::spc* pRW04-$P_{yqeZ}$-*floA-gfp* (*cm*) | This study | RW28+RW394 |
| RW405 | PY79 | Δ*floA::mls* Δ*floT::spc* pRW04-$P_{yuaF}$-*floT-gfp* (*cm*) | This study | RW28+RW392 |
| RW406 | PY79 | Δ*floA::mls* Δ*floT::spc* pRW04-$P_{yqeZ}$-*floA*$_{Nt}$*T*$_{Ct}$-*gfp* (*cm*) | This study | RW28+RW395 |
| RW407 | PY79 | Δ*floA::mls* Δ*floT::spc* pRW04-$P_{yuaF}$-*floT*$_{Nt}$*A*$_{Ct}$-*gfp* (*cm*) | This study | RW28+RW393 |
| RW408 | PY79 | Δ*floA::mls* Δ*floT::spc* pRW04 (*cm*) | This study | RW28+RW201 |
| RW433 | PY79 | Δ*amyE*::$P_{xyl}$-*gfp* (*spc*) | This study | RW3+RW430 |
| RW434 | PY79 | Δ*amyE*::$P_{xyl}$-*gfp* (*spc*) pRW01-$P_{yqeZ}$-*floA-mCherry* (*ery*) | This study | RW3+RW420 |
| RW435 | PY79 | Δ*amyE*::$P_{xyl}$-*gfp* (*spc*) pRW01-$P_{yuaF}$-*floT-mCherry* (*ery*) | This study | RW3+RW421 |
| RW445 | 168 | Δ*trpC2* Δ*pbpC*::pSG5045($P_{xyl}$-*gfp-pbpC*)*cm* | [24] | |
| RW445b | PY79 | Δ*pbpC*::pSG5045($P_{xyl}$-*gfp-pbpC*)*cm* | This study | RW3+RW445 |
| RW554 | PY79 | Δ*lacA*::$P_{yqeZ}$-*floA-mCherry* (*mls*) Δ*pbpC*::pSG5045($P_{xyl}$-*gfp-pbpC*)*cm* | This study | RW45+RW445 (SPP1) |
| RW555 | PY79 | Δ*lacA*::$P_{yuaF}$-*floT-mCherry* (*mls*) Δ*pbpC*::pSG5045($P_{xyl}$-*gfp-pbpC*)*cm* | This study | RW48+RW445 (SPP1) |
| RW312 | PY79 | Δ*pbpC*::*km* | This study | Primers E |
| RW307 | PY79 | Δ*pbpC*::*km* Δ*lacA*::$P_{yqeZ}$-*floA-gfp* (*mls*) | This study | RW312+RW73 |
| RW299 | PY79 | Δ*pbpC*::*km* Δ*lacA*::$P_{yuaG}$-*floT-gfp* (*mls*) | This study | RW312+RW114 |
| RW482 | PY79 | Δ*floA*::*spc* Δ*pbpC*::pSG5045($P_{xyl}$-*gfp-pbpC*)*cm* | This study | RW445b+DL1237 |
| RW488 | PY79 | Δ*floT*::*tet* Δ*pbpC*::pSG5045($P_{xyl}$-*gfp-pbpC*)*cm* | This study | RW330+RW445 (SPP1) |
| RW498 | PY79 | Δ*amyE*::$P_{xyl}$-*gfp-dltD* (*spc*) | This study | RW3+RW494 |
| RW556 | PY79 | Δ*amyE*::$P_{xyl}$-*gfp-dltD* (*spc*) Δ*lacA*::$P_{yqeZ}$-*floA-mCherry* (*mls*) | This study | RW498+RW73 |
| RW557 | PY79 | Δ*amyE*::$P_{xyl}$-*gfp-dltD* (*spc*) Δ*lacA*::$P_{yuaF}$-*floT-mCherry* (*mls*) | This study | RW498+RW114 |
| DL469 | 168 | Δ*dtl*::*tet* | [25] | |
| RW124 | PY79 | Δ*dtl*::*tet* | This study | RW3+DL469 |
| RW104 | PY79 | Δ*dlt*::*tet* Δ*lacA*::$P_{yqeZ}$-*floA-gfp* (*mls*) | This study | RW124+RW73 |
| RW105 | PY79 | Δ*dlt*::*tet* Δ*lacA*::$P_{yuaF}$-*floT-gfp* (*mls*) | This study | RW124+RW114 |
| RW514 | PY79 | Δ*floA*::*cm* Δ*amyE*::$P_{xyl}$-*gfp-dltD* (*spc*) | This study | RW329+RW494 |
| RW513 | PY79 | Δ*floT*::*tet* Δ*amyE*::$P_{xyl}$-*gfp-dltD* (*spc*) | This study | RW330+RW494 |
| RW569 | PY79 | Δ*dtlA*::*tet* | This study | Primers F |
| RW568 | PY79 | Δ*dltA*::*tet* Δ*lacA*::$P_{yuaF}$-*floT-gfp* (*mls*) | This study | RW569+RW114 |
| RW38 | PY79 | Δ*amyE*::$P_{xyl}$-*gfp-mreB* (*spc*) | [26] | |
| RW39 | PY79 | Δ*amyE*::$P_{xyl}$-*gfp-mreBH* (*spc*) | [26] | |
| RW40 | PY79 | Δ*amyE*::$P_{xyl}$-*gfp-mbl* (*spc*) | [26] | |
| RW422 | PY79 | Δ*amyE*::$P_{xyl}$-*gfp-mreB* (*spc*) pRW01-$P_{yqeZ}$-*floA-mCherry* (*ery*) | This study | RW38+RW420 |
| RW423 | PY79 | Δ*amyE*::$P_{xyl}$-*gfp-mreB* (*spc*) pRW01-$P_{yuaF}$-*floT-mCherry* (*ery*) | This study | RW38+RW421 |
| RW491 | 168 | Δ*amyE*::$P_{xyl}$-*mRPFruby-mreB* (*spc*) | [27] | |

| Strain | | Genotype[a] | Reference | Construction |
|---|---|---|---|---|
| RW491b | PY79 | Δ*amyE*::P$_{xyl}$-*mRPFruby-mreB* (*spc*) | This study | RW3+RW491 |
| RW552 | PY79 | Δ*amyE*::P$_{xyl}$-*mRFPruby-mreB* (*spc*) Δ*lacA*::P$_{yqfA}$-*floA-gfp* (*mls*) | This study | RW491b+RW73 |
| RW553 | PY79 | Δ*amyE*::P$_{xyl}$-*mRFPruby-mreB* (*spc*) Δ*lacA*::P$_{yuaF}$-*floT-gfp* (*mls*) | This study | RW491b+RW114 |
| RW576 | 168 | Δ*trpC2* Ω*neo3427* Δ*mreB* Δ*mbl*::*cm* Δ*mreBH*::*erm* Ω(*neo*::*spc*)Δ*rsgI* | [28] | |
| RW610 | PY79 | Δ*floA*::*floA-gfp-tet* Ω(*neo*::*spc*)Δ*rsgI* | This study | RW578+RW576 |
| RW606 | PY79 | Δ*floT*::*floT-gfp-tet* Ω(*neo*::*spc*)Δ*rsgI* | This study | RW579+RW576 |
| RW616 | PY79 | Δ*floA*::*floA-gfp-tet* Ω*neo3427* Δ*mreB* Δ*mbl*::*cm* Δ*mreBH*::*erm* Ω(*neo*::*spc*)Δ*rsgI* | This study | RW578+RW576 |
| RW617 | PY79 | Δ*floT*::*floT-gfp-tet* Ω*neo3427* Δ*mreB* Δ*mbl*::*cm* Δ*mreBH*::*erm* Ω(*neo*::*spc*)Δ*rsgI* | This study | RW579+RW576 |
| RW457 | PY79 | Δ*tagU*::*km* | This study | Primers G |
| RW458 | PY79 | Δ*tagU*::*km* Δ*lacA*::P$_{yqeZ}$-*floA-gfp* (*mls*) | This study | RW457+RW73 |
| RW456 | PY79 | Δ*tagU*::*km* Δ*lacA*::P$_{yuaG}$-*floT-gfp* (*mls*) | This study | RW457+RW114 |
| RW113 | PY79 | Δ*ugtP*::*km* | This study | Primers H |
| RW119 | PY79 | Δ*ugtP*::*km* Δ*lacA*::P$_{floA}$-*floA-gfp* (*mls*) | This study | RW113+RW73 |
| RW117 | PY79 | Δ*ugtP*::*km* Δ*lacA*::P$_{floT}$-*floT-gfp* (*mls*) | This study | RW113+RW114 |

[a] Antibiotic resistance is specified in parentheses if necessary, for *E. coli* plasmids first and second resistance correspond to *E. coli* and *B. subtilis*, respectively.

Supplemental Table S4: List of primers used in this study.

| Purpose | Construct | Name | Sequence 5' - 3' |
|---|---|---|---|
| **Replicative plasmid** | pRW derivates | pRW-MCS_for | AAAAGCATGCTAGCAGATCTCCATGGTACCCGGGAGC |
| | | pRW-MCS_rev | AAAAACTAGTGGCGCGCCGGCGCCGATATCGGATCCATATGACG |
| | | km-ApaI_for | AAAAGGGCCCCAGCGAACCATTTGAGGTG |
| | | km-XhoI_rev | AAAACTCGAGCGATACAAATTCCTCGTAGG |
| | | tet-ApaI_for | TTTTGGGCCCTCTTGCAATGGTGCAGGTTG |
| | | tet-XhoI_for | TTTTCTCGAGCTCTCCCAAAGTTGATCCC |
| | | cm-ApaI_for | AAAAGGGCCCGCAATAGTTACCCTTATTATC |
| | | cm-XhoI_rev | AAAACTCGAGCTGGAGCTGTAATATAAAAAC |
| **Fluorescent-labeled strains** | Flotillins | P$_{yqfeZ}$-EcoRI_for | TAATGAATTCGTGAGCAGTCAACTGTC |
| | | P$_{yuaF}$-KpnI_for | AAAAGGTACCCGCAGCAGTCAGCTGC |
| | | P$_{yuaF}$-SalI_for | AAAAGTCGACCGCAGCAGTCAGCTGC |
| | | mCherry-SphI_rev | AAAAGCATGCTTACTTGTACAGCTCGTCCAT |

| Purpose | Construct | Name | Sequence 5' - 3' |
|---|---|---|---|
| | | $P_{xyl}$-SalI_for | TTTTGTCGACTTTATTGCAATAACAGGT GCTTAC |
| | | gfp-SalI_rev | TTTTGTCGACGTTATTTGTATAGTTC |
| | | gfp-SphI_rev | AAAAGCATGCTTATTTGTATAGTTCATC CATGC |
| | Δ*floA*::*floA-gfp-tet* | floA_for | ATGGATCCGTCAACACTTATG |
| | | gfp-tet_rev | CACATTTCACCCTCCAATAATGTTATTT GTATAGTTCATCCATG |
| | Primers A | tet-gfp_for | CATGGATGAACTATACAAATAACATTAT TGGAGGGTGAAATGTG |
| | | tet-yqfB_rev | CGTTCTCCCTTCTTAGAGAGATTAGAA ATCCCTTTGAGAATG |
| | | yqfB-tet_for | CATTCTCAAAGGGATTTCTAATCTCTCT AAGAAGGGAGAACG |
| | | yqfB_rev | AAGGCATGTACATCCTGAAGC |
| | Δ*floT*::*floT-gfp-tet* | floT_for | ATGACAATGCCGATTATAATG |
| | | tet-yuaH_rev | GGTTCTGCCCTTTCCTTACTCTTAGAAA TCCCTTTGAGAATG |
| | Primers B | yuaH-tet_for | CATTCTCAAAGGGATTTCTAAGAGTAA GGAAAGGGCAGAACC |
| | | yuaH_rev | CAAAGCAGGTCTTACTACAGG |
| | DltD | dltD-XhoI_for | AAAACTCGAGATGAAAAAGCGTTTTTTC GG |
| | | dltD-EcoRI_rev | AAAAGAATTCGGATGAAGTGACTTTTC CGG |
| **Deletions** | Δ*floA*::*cm* | FloA-clean-U-SalI_for | AAAAGTCGACTAGTCGATTGGTGTATT CG |
| | Primers C | FloA-up-cm_rev | GATAATAAGGGTAACTATTGCTCAAGA ATAATATGGTCTGC |
| | | FloA-down-cm_for | GTTTTTATATTACAGCTCCAGCGTATGG TACAGGCAAGA |
| | | FloA-down-BamHI_rev | AAAAGGATCCTTTCGGGCGACATCATT AA |
| | | cm-floA-up_for | GCAGACCATATTATTCTTGAGCAATAGT TACCCTTATTATC |
| | | cm-floA-down_rev | TCTTGCCTGTACCATACGCTGGAGCTG TAATATAAAAAC |
| | Δ*floT*::*tet* | FloT-clean-up-SalI_for | AAAAGTCGACCGGCTTTCGTCCGCCA |
| | Primers D | FloT-up-tet_rev | GAGAACAACCTGCACCATTGCAAGACA GAGGCAATACAAGTTCA |
| | | FloT-down-tet_for | GGGATCAACTTTGGGAGAGAGTTCGCA AAAGGAGCGGAGTTT |
| | | FloT-clean-down-BamHI_rev | AAAAGGATCCGCTGAGAGTGAGCGGT T |
| | | tet-floT-up_for | TGAACTTGTATTGCCTCTGTCTTGCAAT GGTGCAGGTTGTTCTC |
| | | tet-floT-down_rev | AAACTCCGCTCCTTTTGCGAACTCTCT CCCAAAGTTGATCCC |
| | Δ*pbpC*::*km* | pbpC-up-XhoI_for | AAAACTCGAGTGCGGTTATCATTATTAT ACTGG |
| | Primers E | pbpC-up-km_rev | CACCTCAAATGGTTCGCTGATGACTTT CCCCTGCCTTCC |
| | | pbpC-down-km_for | CCTACGAGGAATTTGTATCGCGTTGAG AAAGCGAAAAAGC |
| | | pbpC-down-EcoRI_rev | TTTTGAATTCCTTCTCAGGCAAATATGA TTCC |

| Purpose | Construct | Name | Sequence 5' - 3' |
|---|---|---|---|
| | | km-pbpC_for | GGAAGGCAGGGGAAAGTCATCAGCGAACCATTTGAGGTG |
| | | km-pbpC_rev | GCTTTTTCGCTTTCTCAACGCGATACAAATTCCTCGTAGG |
| | Δ*dltA*::*tet* | dltA-up_for | ATAAGTTCGTCGCGATCTGG |
| | Primers F | dltA-up-tet_rev | GATTGTGAATAGGATGTATTCACCATAGTTATTCTCTCTCCAATTAG |
| | | tet-dltA-up_for | CTAATTGGAGAGAGAATAACTATGGTGAATACATCCTATTCACAATC |
| | | tet-dltA-down_rev | TCATACAAGAACCTCTTCGCCTTAGAAATCCCTTTGAGAATG |
| | | dltA-down-tet_for | CATTCTCAAAGGGATTTCTAAGGCGAAGAGGTTCTTGTATGA |
| | | dltA-down_rev | AGACATATGCCAGCGATTC |
| | Δ*tagU*::*km* | tagU-up-BamHI_for | AAAAGGATCCCGCAGTTTCGTATCGTGAAGC |
| | Primers G | tagU-up-km_rev | CACCTCAAATGGTTCGCTGCGTTTCTCATCCTTTGCACC |
| | | km-tagU-up_for | GGTGCAAAGGATGAGAAACGCAGCGAACCATTTGAGGTG |
| | | km-tagU-down_rev | CCGGATTCATTTACAGGCAAATCGATACAAATTCCTCGTAGG |
| | | tagU-down-km_for | CCTACGAGGAATTTGTATCGATTTGCCTGTAAATGAATCCGG |
| | | tagU-down-EcoRI_rev | TTTTGAATTCGTTTCCGGAAGAGCTCAATCG |
| | Δ*ugtP*::*km* | ugtP-BamHI_for | AAAAGGATCCCTGCGAGAGAACACCTTG |
| | Primers H | ugtP-km_rev | CCTATCACCTCAAATGGTTCGCTGTTAGAAACTGTTACATGCTG |
| | | km-ugtP_for | CAGCATGTAACAGTTTCTAACAGCGAACCATTTGAGGTGATAGG |
| | | km-ugtP_rev | CACTTCAGAGGAGTTTGCTCGATACAAATTCCTCGTAGGCGCTCGG |
| | | ugtP-km_for | CCGAGCGCCTACGAGGAATTTGTATCGAGCAAACTCCTCTGAAGTG |
| | | ugtP-SalI_rev | TTTTGTCGACATCCGTAAAGCCGGTCTG |

### Supplemental Movies

**Supplemental Movie 1: Mobility of FloA is faster than FloT.** Time-lapse fluorescence microscopy of FloA-GFP (top) and FloT-GFP (bottom). The mobility of flotillins (left) and an overlay of their trajectories (right) is shown. Images were taken every 300 ms over 9 s.

**Supplemental Movie 2: Membrane disturbance does not alter flotillin mobility.** Time-lapse fluorescence microscopy of FloA-GFP (top) and FloT-GFP (bottom) after disturbance of membrane properties. Images were taken every 300 ms over 9 s. BNZ=benzyl alcohol, NIS=nisin, VAL=valinomycin.

**Supplemental Movie 3: The C-terminus determines flotillin mobility.** Time-lapse fluorescence microscopy of FloT$_{nt}$A$_{ct}$-GFP (left) and FloA$_{nt}$T$_{ct}$-GFP (right). Images were taken every 300 ms over 9 s.

**Supplemental Movie 4: Flotillin mobility in changes in mutant backgrounds.** Time-lapse fluorescence microscopy of FloA-GFP (top) and FloT-GFP (bottom) in WT, Δ*pbpC,* Δ*dltA-E* and Δ*mreB* Δ*mreBH* Δ*mbl* mutant backgrounds. Images were taken every 300 ms over 9 s.

**Supplemental Movie 5: The mobility of flotillin interaction partners PBP3 and DltD depend on flotillin.** Time-lapse TIRF microscopy of GFP-PBP3 (top) and GFP-DltD (bottom) in WT, Δ*floA* and Δ*floT* backgrounds. Images were taken every 300 ms over 9 s.

**Supplemental Movie 6: Alterations in the cell wall influence flotillin mobility.** Time-lapse fluorescence microscopy of FloA-GFP (top) and FloT-GFP (bottom) after disturbance of the cellular integrity by affecting the cell wall by inhibition of cell wall synthesis and by the formation of protoplasts. Images were taken every 300 ms over 9 s. FOS=fosfomycin, TUN=tunicamycin, AMP=ampicillin, VAN=vancomycin.

**Supplemental Movie 7: The mobility of FloA and FloT imaged with TIRF microscopy.** Time-lapse TIRF microscopy of FloA-GFP (top) and FloT-GFP (bottom). The mobility of flotillins (left) and an overlay of their trajectories (right) is shown. Images were taken every 200 ms over 20 s.

**Supplemental Movie 8: The mobility of MreB is increased in protoplasts.** Time-lapse TIRF microscopy of GFP-MreB cells and protoplasts. Images were taken every 1 s for 100 s to monitor MreB mobility in cells, and every 300 ms over 60 s to monitor MreB mobility in protoplasts.
